# Neural mechanisms of distributed value representations and learning strategies

**DOI:** 10.1101/2021.04.02.438203

**Authors:** Shiva Farashahi, Alireza Soltani

**Author notes:** Corresponding authors: AS, Department of Psychological and Brain Sciences, Dartmouth College, Hanover NH 03755,; SF, Flatiron institute, Simons Foundation, New York NY 10010.

## Abstract

Learning appropriate representations of the reward environment is extremely challenging in the real world where there are many options to learn about and these options have many attributes or features. Despite existence of alternative solutions for this challenge, neural mechanisms underlying emergence and adoption of value representations and learning strategies remain unknown. To address this, we measured learning and choice during a novel multi-dimensional probabilistic learning task in humans and trained recurrent neural networks (RNNs) to capture our experimental observations. We found that participants estimate stimulus-outcome associations by learning and combining estimates of reward probabilities associated with the informative feature followed by those of informative conjunctions. Through analyzing representations, connectivity, and lesioning of the RNNs, we demonstrate this mixed learning strategy relies on a distributed neural code and distinct contributions of inhibitory and excitatory neurons. Together, our results reveal neural mechanisms underlying emergence of complex learning strategies in naturalistic settings.

## Introduction

Successful value-based decision making and learning depend on the ability of the brain to encode and represent relevant information and to update those representations based on feedback in the environment. For example, to be able to learn from an unpleasant reaction to consuming a multi-ingredient meal requires having representations for reward value of certain individual ingredients or combinations of ingredients that were predictive of the outcome (informative attributes). Learning such informative attributes and associated value representations is challenging because feedback is non-specific and scarce (e.g., stomachache after a meal with combinations of ingredients that may never recur), and thus, it is unclear what features or combinations of features are important for predicting the outcomes and must be learned. Such learning becomes even more challenging in high-dimensional environments where the set of possible stimuli or choice options grows exponentially as the number of features or feature instances increases (curse of dimensionality).

Recent studies have shown that human and non-human primates can overcome the curse of dimensionality by learning and incorporating the structure of reward environment to adopt an appropriate learning strategy (Braun et al., 2010; Farashahi, Rowe, et al., 2017; Gershman & Niv, 2010; Leong et al., 2017; Niv et al., 2015; Wilson & Niv, 2012; Wunderlich et al., 2011). For example, when the environment follows a generalizable set of rules such that the values of options can be inferred from their features, humans follow a feature-based learning to estimate reward value of options based on their features instead of learning about individual options (object-based learning) directly (Braun et al., 2010; Dayan & Berridge, 2014; Farashahi, Rowe, et al., 2017; Gershman & Niv, 2010; Leong et al., 2017; Niv et al., 2015; Oemisch et al., 2019; Wilson & Niv, 2012; Wunderlich et al., 2011). In contrast, lack of generalizable rules influence learning by biasing behavior away from fast but imprecise feature-based learning to slower but more precise object-based learning (Farashahi, Rowe, et al., 2017). Despite evidence for adoption of such simple feature-based learning, it is currently unknown whether and how more complex value representations and learning strategies emerge.

Multiple reinforcement learning (RL) and Bayesian models have been proposed to explain how animals can learn informative representations at the behavioral level (Farashahi, Rowe, et al., 2017; Lake et al., 2015, 2017; Leong et al., 2017; Niv et al., 2015; Oemisch et al., 2019; Radulescu et al., 2019). However, all these models assume the existence of certain representations or generative models. For example, there are different proposals for how representations are adopted in connectionist models ranging from extreme localist theories, with single processing unit representations (i.e., grandmother cells) (Barlow, 1972; Parker & Newsome, 1998; Quiroga et al., 2005), to distributed theories with representations that are defined as patterns of activity across a number of processing units (Decharms & Zador, 2000; Hinton, 1986; Hinton et al., 1986). Although local representations are easy to interpret and are the closest to naïve reinforcement learning models, the scarcity of reward feedback and large number of options in high-dimensional environments make these models unappealing. In contrast, distributed representations allow for more flexibility, making them plausible candidates for learning appropriate representations in highdimensional reward environments. Nonetheless, it is currently unknown how multiple representations of reward value and learning strategies emerge over time and what the underlying neural mechanisms are.

To investigate whether and how appropriate representations of reward value and learning strategies are acquired, we examined learning and choice behavior during a novel naturalistic task with a multi-dimensional reward environment and partial generalizable rules in humans. We hypothesized that participants start by learning a simplified representation of the environment before learning more complex representations involving certain combinations or conjunctions of features. This conjunction-based learning would provide an intermediate learning strategy that is faster than object-based learning and more precise than feature-based learning. At the neural level, we hypothesized that such mixed feature- and conjunction-based learning relies on distributed representations of reward value. To test this hypothesis and explore underlying computations and neural mechanisms, we used recurrent neural networks (RNNs) that have been successfully used to address a wide range of neuroscientific questions (Carnevale et al., 2015; Eliasmith et al., 2012; Goudar & Buonomano, 2018; Mante et al., 2013; Masse et al., 2019; Rajan et al., 2016; Song et al., 2016, 2017; Sussillo & Abbott, 2009; Wang et al., 2018; Yang et al., 2019; Zipser & Andersen, 1988). Specifically, we constructed and trained biophysically-plausible recurrent networks of point neurons endowed with plausible reward-dependent Hebbian learning rules (Gerstner et al., 2014; Loewenstein & Seung, 2006; Pfeiffer et al., 2010) to perform our learning task. We then applied a combination of representational similarity analysis (Kriegeskorte et al., 2008), connectivity pattern analysis, and lesioning of the trained RNNs to reveal neural mechanisms underlying the evolution of value representations and learning strategies observed in our experimental paradigm.

## Results

### Learning about informative features and conjunctions of features in multi-dimensional environments

Building upon our previous work on feature-based vs. object-based learning (Farashahi, Rowe, et al., 2017), we designed a novel multi-dimensional probabilistic learning task (mdPL) that allows study of the emergence of intermediate learning strategies. In this task, human participants learned stimulus-outcome associations by selecting between pairs of targets defined by their visual features (color, pattern, and shape) followed by a binary reward feedback. Moreover, we asked subjects to provide their estimates of reward probabilities for individual objects during five bouts of estimation trials throughout the experiment (see **Methods** for more details). Critically, the reward probability associated with selection of each stimulus was determined by the combination of its features such that one “informative” feature and the conjunctions of the other two “non-informative” features could partially predict reward probabilities (**Supplementary Figure 1**). For example, square-shaped stimuli could be on average more rewarding than diamond-shaped stimuli, whereas stimuli with different colors (or patterns) were equally rewarding on average, corresponding to shape being the informative feature and color and pattern being the non-informative features. At the same time, stimuli with certain combinations of color and pattern (e.g., solid blue) could be more rewarding than stimuli with other combinations of color and pattern, making conjunctions of color and pattern to be the informative conjunction.

Analyzing participants’ performance and choice behavior suggested that the majority of participants understood the task and learned about stimuli while reaching their steady-state performance in about 150 trials. This was evident by calculating the average total harvested reward as well as the probability of choosing the better stimulus (i.e., stimulus with higher probability of reward) in each trial over the course of the experiment (**Figure 1A**). To identify the learning strategy adopted by each participant, we fit individual participants’ choice behavior using 24 different reinforcement learning (RL) models that relied on feature-based, object-based, mixed feature- and conjunction-based, or mixed feature- and object-based learning strategies (see **Methods**). By fitting participants’ choice data and computing AIC and BIC per trial (AICp and BICp; see **Methods**) as different measures of goodness-of-fit (Farashahi et al., 2018, 2020), we found that one of the mixed feature- and conjunction-based models provided the best overall fit for all data and throughout the experiment (**Supplementary Table 1, Figure 1B, Supplementary Figure 2**). In this model (the F+C_1_ model), the decision maker updates the estimated reward probabilities associated with the informative feature and the informative conjunction of the selected stimulus after each feedback while forgetting reward probabilities associated with the unchosen stimulus (by decaying those values toward 0.5). As a result, this mixed feature- and conjunction-based model was able to learn quickly without compromising precision in the estimated reward probabilities (**Supplementary Figure 3**). Consistent with this, we found that the difference in the goodness-of-fit between this model and the second-best model (feature-based model) increased over time (**Figure 1B**), pointing to more use of the mixed learning strategy by the participants.

**Figure 1.**
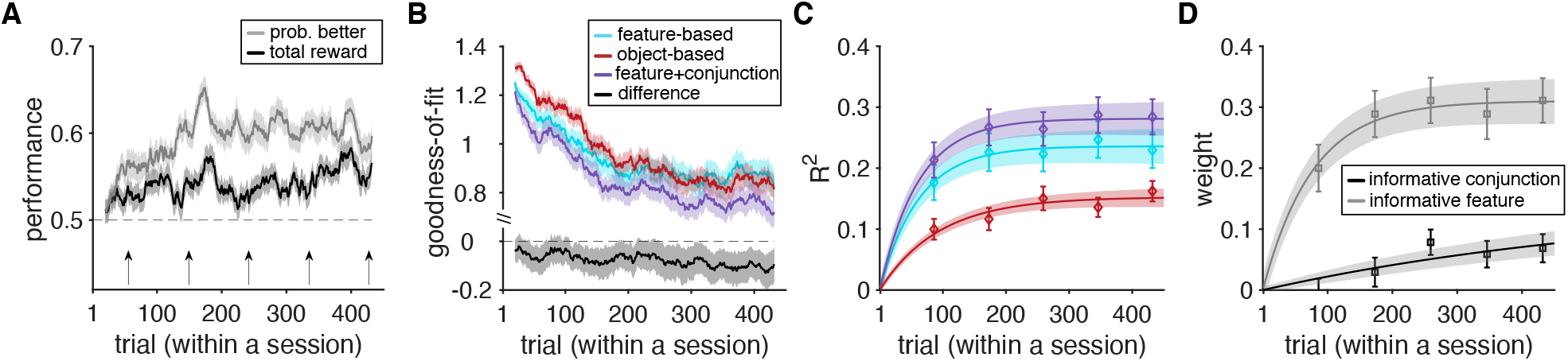
Evidence for adoption of mixed feature- and conjunction-based learning. (**A**) Time course of performance and learning during the experiment. Plotted are the average total harvested reward and probability of selecting the stimulus with higher probability of reward (better option) in a given trial within a session of the experiment. The running average over time is computed using a moving box with the length of 20 trials. Shaded areas indicate s.e.m., and the dashed line shows chance performance. (**B**) Plotted is the goodness-of-fit based on the average AIC per trial, AICp, for the feature-based model, object-based model, and the best mixed feature- and conjunction-based (F+C_1_) model. The smaller value corresponds to a better fit. The black curve shows the difference between the goodness-of-fit for the F+C_1_ and feature-based models. (**C**) The plot shows the time course of explained variance (*R^2^*) in participants’ estimates based on different GLMs. Color conventions are the same as in panel B with cyan, red, and purple curves representing *R^2^* based on the feature-based, object-based, and F+C_1_ models, respectively. The solid line is the average of fitted exponential function to each participant’s data and shaded areas indicate s.e.m. of the fit. (**D**) Time course of adopted learning strategies measured by fitting participants’ estimates of reward probabilities. Plotted is the weight of the informative feature and informative conjunction in the F+C_1_ model. Error bars indicate s.e.m. The solid line is the average of fitted exponential function to each participant’s data, and shaded areas indicate s.e.m. of the fit.

In addition to choice trials, we also examined estimation trials to determine learning strategies adopted by individual participants over the course of the experiment. To that end, we used generalized linear models (GLMs) to fit participants’ estimated reward probabilities associated with each stimulus based on the predicted reward probabilities using different learning strategies (see **Methods**). The explained variance from different GLMs confirmed that the F+C_1_ model captured participants’ estimates the best (**Figure 1C**). In addition, the time course of extracted weights for the informative feature regressor and the informative conjunction regressor suggested that participants learned the informative feature followed by the informative conjunction. More specifically, we fit the time course of extracted weights to estimate the time constant at which these weights reached their asymptotes (see Eq. 3 in **Methods**). We found that the time constant of increase in the weight of informative feature (median±IQR = 76.3±37.6) was an order of magnitude smaller than the time constant of increase in the weight of informative conjunction (median±IQR = 997.3±48.5; two-sided sign-rank test, *P* = 1.08×10^−5^, *d* = 0.67, *N* = 67; **Figure 1D**), indicating much faster learning of the informative feature comparted with the informative conjunction.

We also analyzed data from the excluded participants. However, this analysis did not provide any evidence that excluded participants adopted learning strategies qualitatively different from those used by the remaining participants (**Supplementary Figure 4**). Instead, it showed that excluded subjects simply did not learn the task. Together, these results demonstrate that our participants were able to learn more complex representations of reward value over time and combined information from these representations with simple representation of individual features to increase their accuracy without slowing down learning.

To assess learning strategies more directly and without any model fitting, we also examined participants’ response to different types of reward feedback (reward vs. no reward). To that end, we calculated differential response to reward feedback by computing the difference between the tendency to select a feature (or a conjunction of features) of the stimulus that was selected on the previous trial and was rewarded vs. when it was not rewarded (see **Methods, Supplementary Figure 5**). Our prediction was that participants who learned about the informative feature and informative conjunction, as assumed in the F+C_1_ model, should exhibit positive differential response for both the informative feature and informative conjunction. In contrast, differential response would be positive for the informative feature (and possibly the non-informative features) for participants who only learned about individual features and not their conjunctions. We used our measure of goodness-of-fit to determine whether a participant adopted feature-based or mixed feature- and conjunction-based learning (62 out of 67 subjects). We did not calculate differential response for participants in the small minority of subjects (5 out of 67) who adopted the object-based strategy.

We found that differential response for informative conjunction and informative feature was overall positive for participants who adopted the best mixed feature- and conjunction-based learning strategy (the F+C_1_ model) (two-sided sign-rank test; informative feature: *P* = 8×10^−4^, *d* = 0.59, *N* = 41; informative conjunction: *P* = 0.029, *d* = 0.19, *N* = 41), whereas differential response for non-informative features was not distinguishable from 0 (two-sided sign-rank test; *P* = 0.23, *d* = 0.058, *N* = 41; **Figure 2A, C**). In contrast, for participants who adopted the feature-based learning strategy, only differential response for the informative feature was significantly larger than zero (two-sided sign-rank test; informative feature: *P* = 0.022, *d* = 0.37, *N* = 21; non-informative feature: *P* = 0.09, *d* = 0.25, *N* = 21) with differential response for the informative feature being larger than that of non-informative features (*P* = 0.005, *d* = 0.43, *N* = 21; **Figure 2B, D**). Moreover, for these participants, differential response for the informative conjunction was not distinguishable from zero (two-sided sign-rank test; *P* = 0.27, *d* = 0.17, *N* = 21).

**Figure 2.**
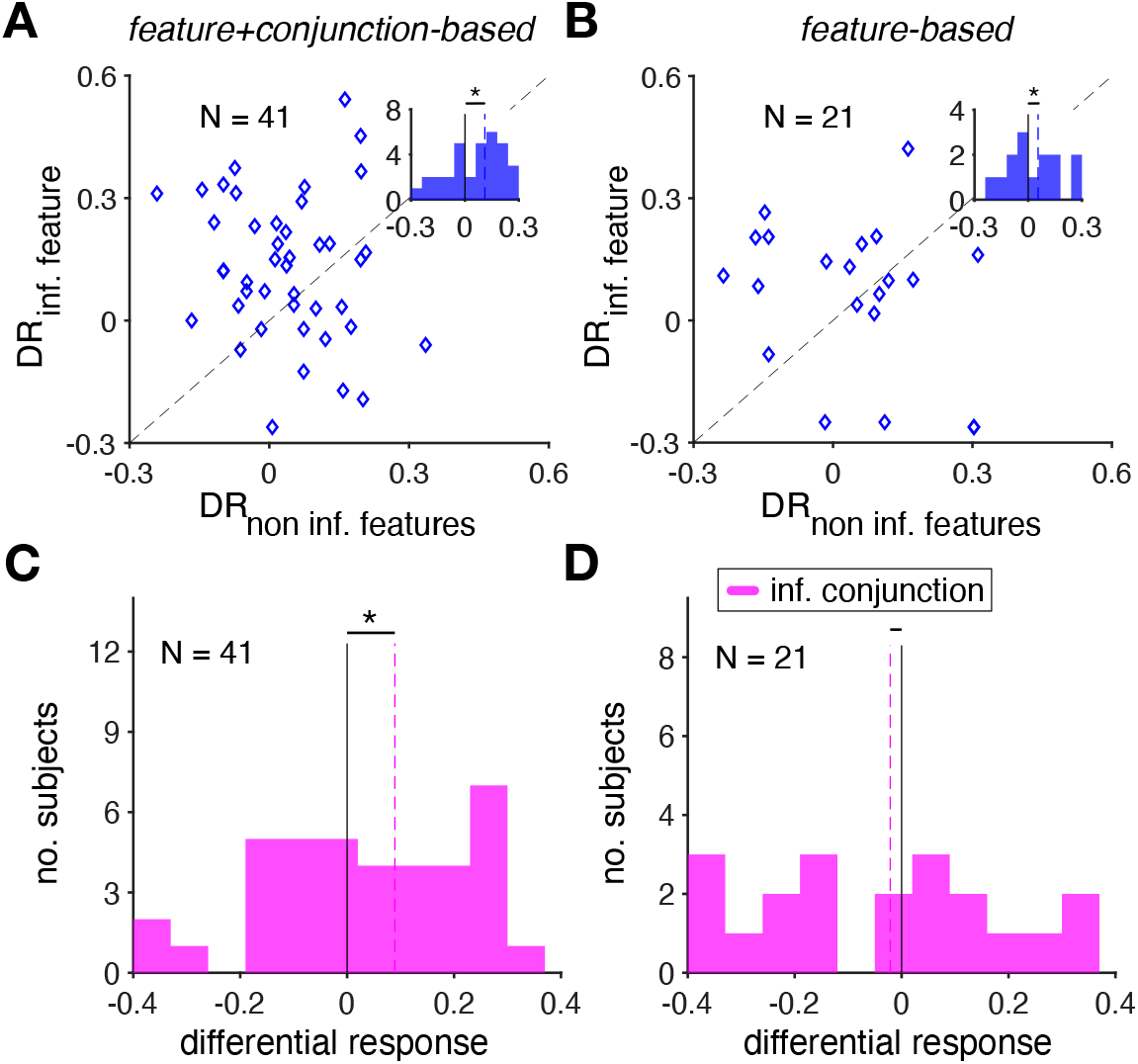
Direct evidence for adoption of mixed feature- and conjunction-based learning strategy. (**A–B**) Plot shows the differential response for the informative vs. non-informative features for participants whose choice behavior was best fit by the F+C_1_ model (A) and for participants whose choice behavior was best fit by the feature-based model (B). The insets show the histogram of difference between differential response of the informative feature and differential response of the non-informative features. The dashed lines show the median values across participants. The significance level of the test is coded as: *P* < 0.05 (*) using two-sided sign-rank test. (**C–D**) Plot shows the histogram of differential response for the informative conjunction for participants whose choice behavior was best fit by the F+C_1_ model (C), and for participants whose choice behavior was best fit by the feature-based model (D). The dashed lines show the median values across participants. The significance level of the test is coded as: *P* < 0.05 (*) using two-sided sign-rank test.

Overall, our experimental results demonstrate for the first time that when learning about high-dimensional stimuli, humans adopt a mixed learning strategy that involves learning about the informative feature as well as the informative conjunction of the non-informative features. Importantly, the informative feature is learned quickly and is slowly followed by learning about the informative conjunction, indicating the gradual emergence of more complex representations over time.

### RNNs exhibit intermediate learning strategies similar to human participants

To account for our experimental observations and gain insight into neural mechanisms underlying the emergence of different value representation and learning strategies, we trained recurrent neural networks (RNNs) of excitatory and inhibitory units (with 4 to 1 ratio) to perform our task in two steps (**Figure 3**). In the first (training) step, we trained RNNs to learn the temporal dynamics of input to output association for estimating reward probabilities of 27 three-dimensional stimuli used in our learning task. In this step, for each training session of 270 trials, reward probabilities were randomly assigned to different stimuli to enable the network to learn a universal solution for learning reward probabilities in multi-dimensional environments with different levels of generalizability, (i.e., how well reward probabilities based on features and conjunctions predict reward probabilities associated with all stimuli). We used the stochastic gradient descent (SGD) method to train the networks (i.e., adjust the connection weights) to perform a general learning task based on a three-factor, reward-dependent Hebbian learning rule (see **Methods** for more details). In the second step (simulation of the experiment), we stopped SGD and simulated the behavior of the trained RNNs in a session of a specific learning task with reward probabilities similar to our experimental paradigm. In this step, only connections endowed with the plasticity mechanism were modulated after receiving reward feedback in each trial. The overall task structure used in these simulations was similar to our experimental paradigm with a simple modification where only one stimulus was shown in each trial and the network had to learn the reward probability associated with that stimulus. Using this approach, we avoided complexity related to decision-making processes and mainly focused on learning aspects of the task. In summary, in the first step, RNNs learned to perform a general multidimensional reward learning task, whereas in the second step, the trained RNNs were used to perform our task.

**Figure 3.**
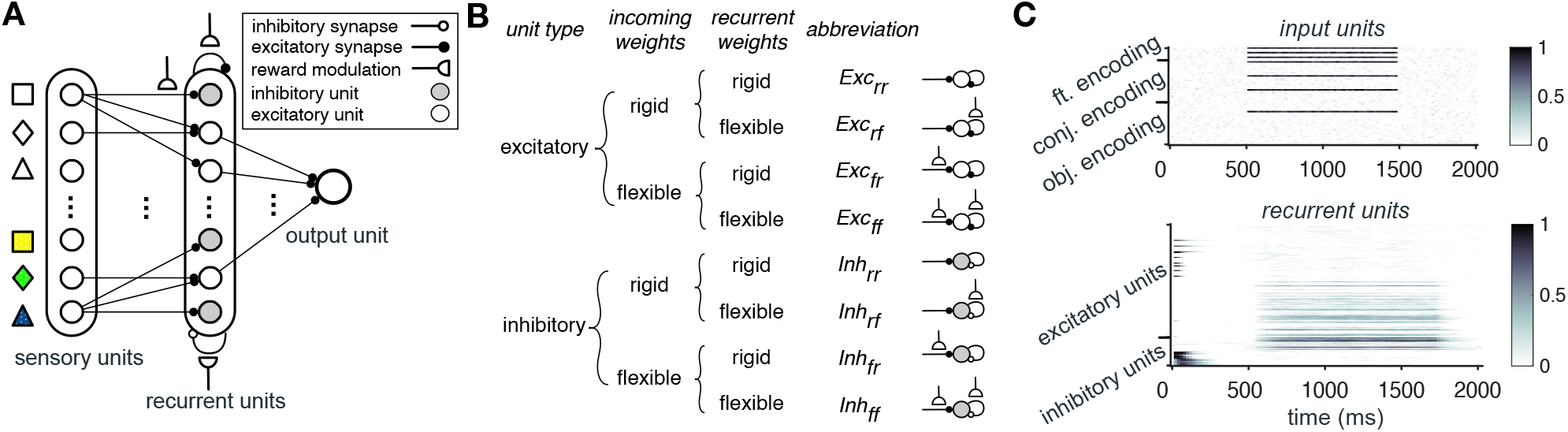
Architecture of the RNN models. (**A**) The models consist of three layers: sensory units, recurrent units, and an output unit. The recurrent units included both excitatory and inhibitory pools that receive input from 63 sensory units encoding individual features (*N* = 9), conjunctions of features (*N* = 27), and object-identity of each stimulus (*N* = 27). Among the recurrent units (*N* = 120) only the excitatory recurrent units project to the output unit. Half of the connections from sensory units and the connections between recurrent units were endowed with reward-dependent plasticity. (**B**) Based on the type of units and presence/absence of reward-dependent plasticity, recurrent units could be grouped into 8 disjoint populations: *Exc_rr_* and *Inh_rr_* corresponding to populations with no plastic sensory or recurrent connections (rigid weights indicated by subscript *r*); *Exc_fr_* and *Inh_fr_* corresponding to populations with plastic sensory input only (flexible weights indicated by subscript *f*); *Exc_rf_* and *Inh_rf_* corresponding to populations with plastic recurrent connections only; and *Exc_ff_* and *Inh_ff_* corresponding to populations with plastic sensory input and plastic recurrent connections. (**C**) Activity of the sensory and recurrent units in an example trial. Upon presentation of a stimulus, three feature-encoding, three conjunction-encoding, and one object-identity encoding units become active and give rise to activity in the excitatory and inhibitory recurrent units.

We first confirmed that networks’ estimate of reward probabilities during the simulation of our task matched that of the human participants (**Figure 4A**). Next, we utilized generalized linear models (GLMs) to fit reward probability estimates in order to identify learning strategies adopted by the RNNs during the course of the experiment. The explained variance of fit of RNNs’ estimates confirmed that similar to our human participants, estimates of RNNs over time were best fit by the F+C_1_ model (**Figure 4B**).

**Figure 4.**
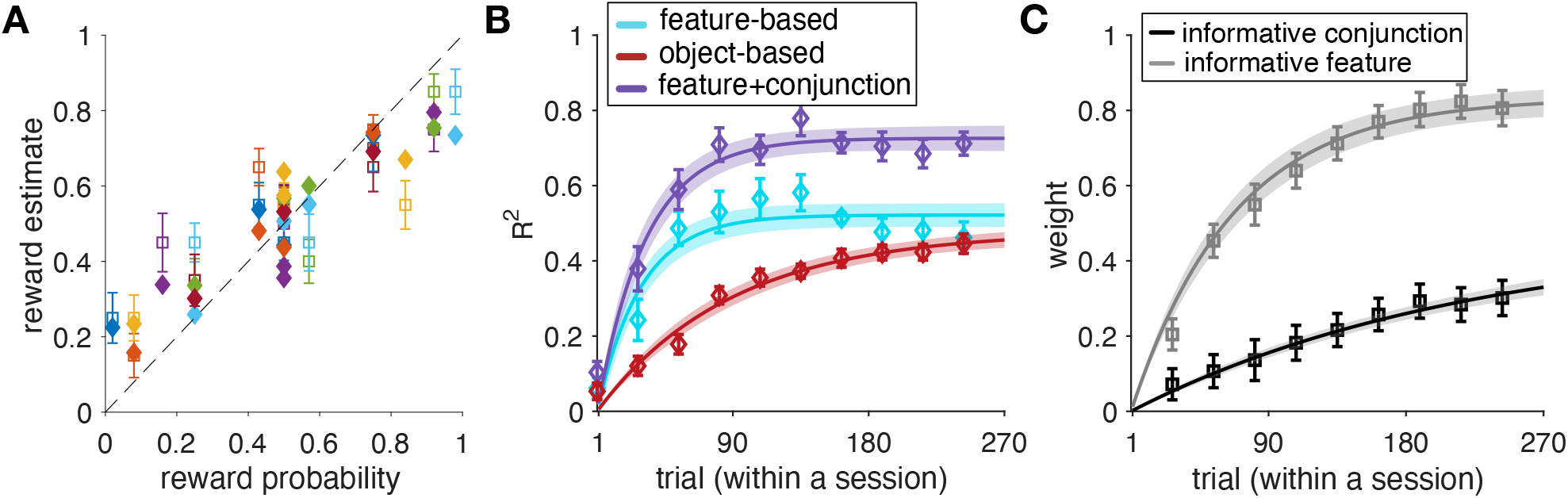
RNNs can capture main behavioral results. (**A**) Plotted are average estimates at the end of a simulated session of our learning task, for 50 instances of the simulated RNNs (filled) and average value of participants’ reward estimate (hollow) vs. actual reward probabilities associated with each stimulus (each symbol represents one stimulus). Error bars represent s.e.m., and the dashed line is the identity. (**B**) The plot shows the time course of explained variance (*R^2^*) in RNNs’ estimates based on different models. Error bars represent s.e.m. The solid line is the average of exponential fits to RNNs’ data, and the shaded areas indicate s.e.m. of the fit. (**C**) Time course of adopted learning strategies measured by fitting the RNNs’ output. Plotted is the weight of the informative feature and informative conjunction in the F+C_1_ model. Error bars represent s.e.m. The solid line is the average of exponential fits to RNNs’ data, and the shaded areas indicate s.e.m. of the fit.

Moreover, the extracted weights for the informative feature and informative conjunction suggest that similar to our human participants, RNNs learn the informative feature followed by the informative conjunction (**Figure 4C**). This was reflected in the time constant of increase in the weight of informative feature (median±IQR = 80.12±68.44) being significantly smaller than that of the informative conjunction (median±IQR = 200.75±107.19; two-sided sign-rank test, *P* = 0.008, *d* = 0.36, *N* = 50).

Together, these results illustrate that our proposed biophysically-plausible RNNs are able to replicate our experimental findings and exhibit transition between learning strategies over the course of the experiment. Next, we examined the response of RNNs’ units to identify the neural substrates underlying different learning strategies and their emergence over time.

### Learning strategies are reflected in the response of different neural types

To investigate value representations that accompany the evolution of learning strategies observed in our task, we applied representational similarity analysis (Kriegeskorte et al., 2008) to the response of units in the trained RNNs. More specifically, we examined how dissimilarity in the response of recurrent units (response dissimilarity matrix) can be predicted based on dissimilarity of reward probabilities calculated according to different learning strategies (reward probability dissimilarity matrices; see **Methods** for more details). To that end, we used GLMs to estimate the normalized weights of the reward probability dissimilarity matrices in predicting the response dissimilarity matrix. Using this method, we were able to quantify how much representations in recurrent units reflect or accompany a particular learning strategy.

We found that recurrent units with plastic sensory input (*Exc_fr_*, *Exc_ff_, Inh_fr_*, and *Inh_ff_*) show strong but contrasting response to reward probabilities associated with stimuli and their features (**Figure 5C, D, G, H**). Importantly, dissimilarity of reward probabilities based on the informative feature could better predict dissimilarity of response in inhibitory populations with plastic sensory input (*Inh_fr_* and *Inh_ff_*; **Figure 5G, H**) compared to the excitatory populations with plastic sensory input (*Exc_fr_* and *Exc_ff_*; **Figure 5C, D**). This was reflected in the difference between the weights of the informative feature for these inhibitory and excitatory populations being significantly larger than 0 (median±IQR = 0.36±0.21; two-sided sign-rank test, *P* = 0.004, *d* = 0.99, *N* = 50).

**Figure 5.**
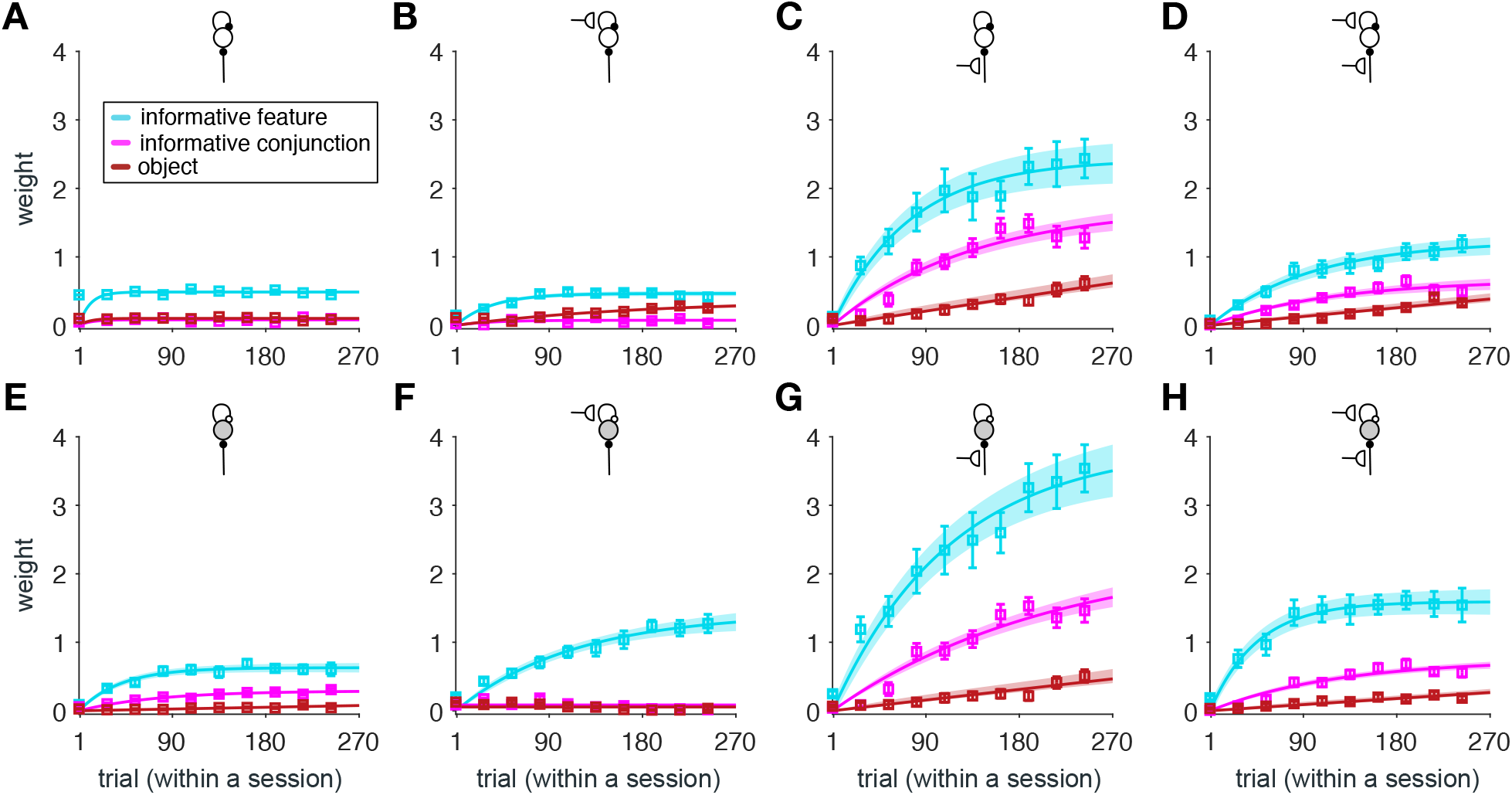
Response of different types of recurrent units show differential degrees of similarity to reward probabilities based on different learning strategies. (**A–D**) Plotted are the estimated weights for predicting the response dissimilarity matrix of different types of recurrent populations (indicated by the inset diagrams explained in Figure 3B) using the dissimilarity of reward probabilities based on the informative feature, informative conjunction, and object. Error bars represent s.e.m. The solid line is the average of fitted exponential functions to RNNs’ data, and the shaded areas indicate s.e.m. of the fit. (**E–H**) Same as A–D but for inhibitory recurrent populations. Dissimilarity of reward probabilities of the informative feature can best predict dissimilarity of response in inhibitory populations, while dissimilarity of reward probabilities of the objects can best predict dissimilarity of response in excitatory populations.

In contrast, dissimilarity of reward probabilities based on objects could better predict dissimilarity of response in excitatory populations with plastic sensory input (*Exc_fr_* and *Exc_ff_*; **Figure 5C, D**) compared to the inhibitory populations with plastic sensory input (*Inh_fr_* and *Inh_ff_*; **Figure 5G, H**). This was reflected in the difference between the weights of the reward dissimilarity matrix based on objects for these excitatory and inhibitory populations being larger than 0 (median±IQR = 0.06±0.07; two-sided sign-rank test, *P* = 0.002, *d* = 1.32, *N* = 50).

Finally, we did not find any significant difference between how dissimilarity of reward probabilities based on the informative conjunction predicts dissimilarity in the response of inhibitory and excitatory populations. The difference between the weights of the informative conjunction for inhibitory and excitatory populations was not significantly different from 0 (median±IQR = −0.004±0.08; two-sided sign-rank test; *P* = 0.56, *d* = 0.45, *N* = 50). However, we found that dissimilarity of reward probabilities based on the informative conjunction can better predict the dissimilarity of response in recurrent units with plastic sensory input only (*Exc_fr_* and *Inh_fr_*) (**Figure 5C, G**) compared to recurrent units with plastic sensory input and plastic recurrent connections (*Exc_ff_* and *Inh_ff_*) (**Figure 5D, H**). This was reflected in the difference between the weights of informative conjunction for these populations being larger than 0 (median±IQR = 0.57±0.17; two-sided sign-rank test; *P* = 0.002, *d* = 1.91, *N* = 50).

Together, these results provide a few novel predictions about the representations of reward value by different neural types. First, only neurons with plastic sensory input exhibit representations of value compatible with the evolution of learning strategies observed in our task. Second, there is an opponency between representations of feature and object values by excitatory and inhibitory neurons; excitatory neurons better represent object values, whereas inhibitory neurons better represent feature values. Finally, neurons with only plastic sensory input exhibit more pronounced representations of conjunction values. To better understand the roles of different neural types in learning, we next examined how the aforementioned representations emerge over time.

### Connectivity pattern reveals distinct contributions of excitatory and inhibitory neurons

To study how different representations emerge in the RNNs, we probed connection weights at the end of the training step and examined how these weights were modulated by reward feedback during the simulation of our task. We refer to the weights at the end of the training step as naïve weights because at that point, the network has not been exposed to the specific relationship between reward probabilities of stimuli and their features used in our task. These naïve weights are important because they reveal the state of connections due to learning in high-dimensional environments with different levels of generalizability (**Supplementary Figure 6**). Moreover, these weights determine the activity of recurrent units that influences subsequent changes in connections due to reward feedback in our task.

We found that after the training in high-dimensional environments with different levels of generalizability, feature-encoding units were connected more strongly to inhibitory populations with plastic sensory input (*Inh_fr_* and *Inh_ff_*; **Figure 6A**). Specifically, the average value of naïve weights from feature-encoding units to inhibitory populations with plastic sensory input was significantly larger than the average value of naïve weights from feature-encoding units to other populations (median±IQR = 0.07±0.02, two-sided sign-rank test; *P* = 5×10^−4^, *d* = 1.21, *N* = 50). In contrast, object-identity encoding units were connected more strongly to the excitatory units with plastic sensory input (*Exc_fr_* and *Exc_ff_*). Specifically, the average value of naïve weights from object-encoding units to excitatory populations with plastic sensory input was significantly larger than the average value of naïve weights from object-encoding units to other populations (median±IQR = 0.06±0.03, two-sided sign-rank test; *P* = 9×10^−3^, *d* = 0.45, *N* = 50). Finally, we did not find any evidence for differential connections between sensory units that encoded conjunctions to different types of recurrent units (median±IQR < 0.006±0.007, two-sided sign-rank test; *P* > 0.13, *d* = 0.10, *N* = 50).

**Figure 6.**
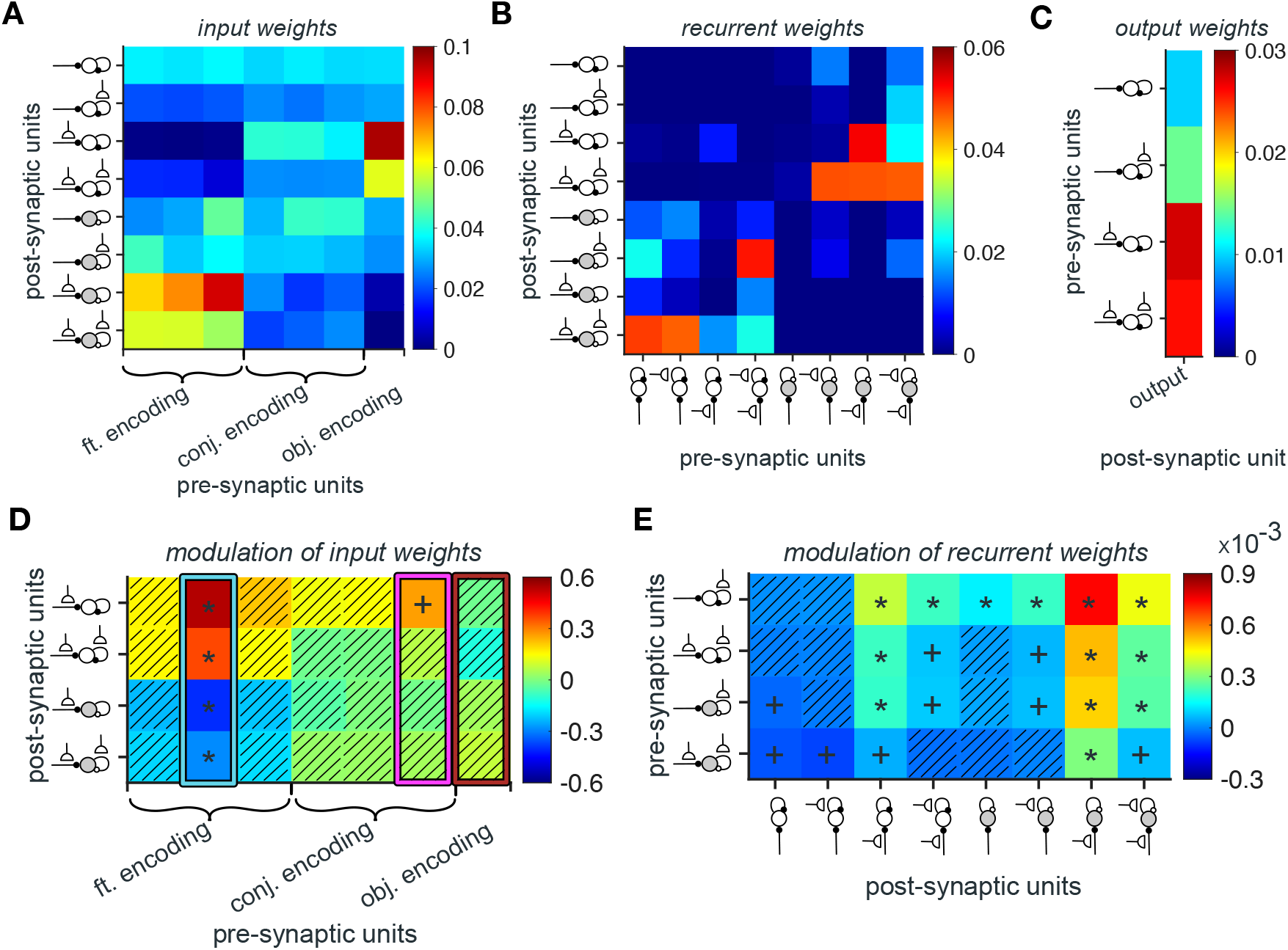
RNNs’ naïve weights at the end of the training step and their subsequent rates of change due to reward feedback during the simulation of our task. (**A–C**) Plotted is the average strength of the naïve weights from feature-encoding, conjunction-encoding, and object-identity encoding units to eight types of recurrent units (A), naïve weights between eight types of recurrent units (B), and naïve weights from eight types of recurrent units to the output unit (C). (**D**) Plotted is the average rate of value-dependent changes in the connection weights from feature-encoding, conjunction-encoding, and object-identity encoding units to recurrent units with plastic sensory input, during the simulation of our task. Asterisks and plus sign indicate two-sided and one-sided significant rates of change (*P*<0.05), respectively. Hatched squares indicate connections with rates of change that were not significantly different from zero (*P*>0.05). Highlighted rectangles in cyan, magenta, and red indicate the values for input from sensory units encoding the informative feature, the informative conjunction, and object-identity, respectively. (**E**) Plotted are the average rates of change in connection weights between recurrent units. Conventions are the same as in D.

With regards to recurrent connections, we found stronger connections between excitatory and inhibitory units than self-excitation and self-inhibition. Specifically, the average value of naïve weights from excitatory to inhibitory populations and vice versa were significantly larger than the average value of naïve excitatory-excitatory and inhibitory-inhibitory weights (median±IQR = 0.017±0.009, two-sided sign-rank test; *P* < 10^−10^, *d* = 0.41, *N* = 50; **Figure 6B**). Among these weights, we found that the average naïve weights from inhibitory populations with plastic sensory input (*Inh_fr_* and *Inh_ff_*) to excitatory populations with plastic sensory input (*Exc_fr_* and *Exc_ff_*) was significantly larger than the average naïve weights between other excitatory and inhibitory populations (median±IQR = 0.036±0.006, two-sided sign-rank test; *P* < 10^−10^, *d* = 1.31, *N* = 50). Similarly, the average value of naïve weights from excitatory populations with no plastic sensory input (*Exc_rr_* and *Exc_rf_*) to inhibitory population with plastic sensory input and plastic recurrent connections (*Inh_ff_*) was significantly larger than the average naïve weights between other excitatory and inhibitory recurrent populations (median±IQR = 0.041±0.005, two-sided sign-rank test; *P* < 10^−10^, *d* = 1.7, *N* = 50).

Finally, among excitatory pools, those with plastic sensory input had the strongest influence on the output unit. This was reflected in the naïve weights from these populations to the output unit being significantly larger than the average value of naïve weights from excitatory population with no plastic sensory input to the output unit (median±IQR = 0.015±0.008, two-sided sign-rank test; *P* = 0.005, *d* = 0.51, *N* = 50; **Figure 6C**).

Together, analyses of naïve weights illustrate that learning in environments with a wide range of generalizability results in a specific pattern of connections. More specifically, feature-encoding sensory units are more strongly connected to inhibitory neurons, whereas object-encoding sensory units are more strongly connected to excitatory neurons. Moreover, stronger cross connections between inhibitory and excitatory neurons indicate an opponency between the two neural types in shaping complex learning strategies.

Due to activity-dependence of the learning rule, stronger connections could be modified more dramatically during the simulation of our task, and thus, are more crucial for the observed behavior. To directly test this, we used GLMs to fit plastic input weights from sensory units during the course of learning in our task. More specifically, we used reward probabilities associated with different aspects of the presented stimulus (features, conjunctions of features, and object-identity) to predict connection weights from sensory to recurrent units with plastic sensory input. This allowed us to measure the rates of change in connection weights due to learning in our task and thus, the contributions of different units to observed learning behavior. We also measured changes in plastic connections between recurrent units over time (see **Methods** for more details).

We found that connection weights from feature-encoding units to inhibitory populations showed a value-dependent reduction as the result of learning in our task, which was only significant in the connections from sensory units encoding the informative feature (median±IQR = −0.34±0.06, twosided sign-rank test; *P* < 0.01, *d* > 0.44, *N* = 50; **Figure 6D**). At the same time, the connection weights from feature-encoding units to excitatory populations showed a significant value-dependent increase (median±IQR = 0.45±0.07, two-sided sign-rank test; *P* < 0.01, *d* > 0.37, *N* = 50). The connection weights from sensory units encoding the informative conjunction to excitatory population with plastic sensory input showed a trend for value-dependent increase (median±IQR = 0.26±0.06, two-sided sign-rank test; *P* = 0.08, *d* = 0.18, *N* = 50).

Finally, analysis of plastic recurrent connections over time showed that the rates of change in weights to the inhibitory population with plastic sensory input (*Inh_fr_* and *Inh_ff_*) was modulated more strongly than all other recurrent connections (median±IQR = 3.8×10^−4^±1.8×10^−5^, two-sided sign-rank test; *P* = 2.8×10^−4^, *d* = 0.35, *N* = 50; **Figure 6E**). Among connections to the these populations, the rates of change in weights from excitatory population with plastic recurrent connections (*Exc_rf_* and *Exc_ff_*) was stronger compared to the rates of change in weights from inhibitory population with plastic sensory input (*Inh_rf_* and *Inh_ff_*) (median±IQR = 2.1×10^−4^±1.4×10^−5^, two-sided sign-rank test; *P* = 0.01, *d* = 0.20, *N* = 50). These results suggest the importance of inhibitory population with plastic sensory input for the observed changes in learning strategies over time.

Together, our results on changes in connectivity pattern due to learning in our task illustrate that learning about stimuli leads to simultaneous increase in connection strength from feature- and conjunction-encoding units to excitatory populations and a decrease in connection strength from feature-encoding units to inhibitory populations. These simultaneous changes effectively result in excitation and disinhibition of excitatory populations, suggesting a role for a delicate interplay between inhibition and excitation in acquiring and adopting mixed feature- and conjunction-based learning strategies.

### Causal role of certain connections in emergence of learning strategies

As mentioned above, we found strong connections between the inhibitory and excitatory populations with plastic sensory input (**Figure 6B, E**). This suggests that these connections could play an important role in the emergence of mixed feature- and conjunction-based learning strategies over time. To test this, in two separate sets of simulations we lesioned connections from the inhibitory populations with plastic sensory input (*Inh_fr_* and *Inh_ff_*) to the excitatory populations with plastic sensory input (*Exc_fr_* and *Exc_ff_*) and vice versa. We found that the network with lesioned connections from the inhibitory populations with plastic sensory input to the excitatory populations with plastic sensory input exhibited an object-based learning strategy (**Figure 7A**). Consistent with this result, unlike the intact network, there was no value-dependent changes in connection weights from feature- and conjunction-encoding units to excitatory and inhibitory populations (**Figure 7B**). However, we found significant value-dependent changes in connection weights from object-encoding units to recurrent units with plastic sensory input and recurrent connections (median±IQR = 0.12±0.04, twosided sign-rank test; *P* = 0.04, *N* = 50), pointing to the role of excitatory neurons in driving object-based learning.

**Figure 7.**
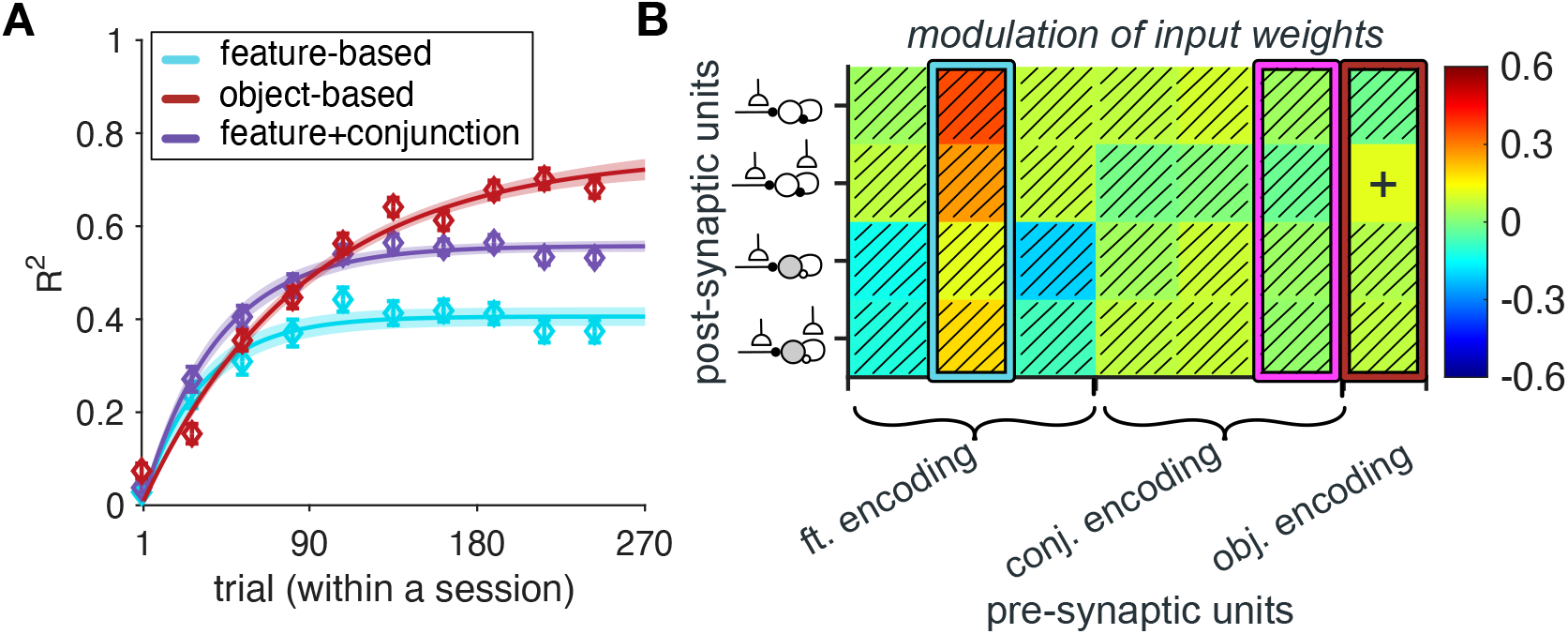
Lesioning recurrent connections from inhibitory populations with plastic sensory input to excitatory populations with plastic sensory input results in drastic changes in the behavior of the RNNs. (**A**) The plot shows the time course of explained variance (*R^2^*) in RNNs’ estimates based on different models. Error bars represent s.e.m. The solid line is the average of exponential fits to RNNs’ data, and the shaded areas indicate s.e.m. of the fit. (**B**) Plotted is the average rate of value-dependent changes in the connection weights from feature-encoding, conjunction-encoding, and object-identity encoding units to recurrent units with plastic sensory input, during the simulation of our task. Plus sign indicates one-sided significant rates of change (*P*<0.05), and hatched squares indicate connections with rates of change that were not significantly different from zero (*P*>0.05). Highlighted rectangles in cyan, magenta, and red indicate the values for input from sensory units encoding the informative feature, the informative conjunction, and object-identity, respectively.

Consistent with these observations, we also found that the dissimilarity of the stimulus reward probabilities in the network with lesioned connections from the inhibitory populations with plastic sensory input to the excitatory populations with plastic sensory input was best predicted by the dissimilarity of response based on objects (**Supplementary Figure 7**). This was reflected in the difference between the weight of object and the weights of informative feature and informative conjunction for this population being larger than 0 (median±IQR = 0.06±0.12, two-sided sign-rank test; *P* = 5.27×10^−4^, *d* = 0.37, *N* = 50).

In contrast, lesioning the connections from the excitatory populations with plastic sensory input (*Exc_fr_* and *Exc_ff_*) to the inhibitory populations with plastic sensory input (*Inh_fr_* and *Inh_ff_*) did not strongly impact the emergence of mixed feature- and conjunction-based learning (**Supplementary Figure 8A, B**). However, in this lesioned network, only the connection weights from feature-encoding units to excitatory populations showed a value-dependent increase over time (median±IQR = −0.34±0.06, two-sided sign-rank test; *P* < 0.01, *N* = 50, **Supplementary Figure 8C**). In addition, we found a decrease in the explanatory power of dissimilarity of the informative feature and the informative conjunction in predicting the dissimilarity of the stimulus reward probabilities compared to the intact network (median±IQR = −0.17±0.29, two-sided sign-rank test; *P* = 2.65×10^−5^, *d* = 0.36, *N* = 50; **Supplementary Figure 9**). However, the informative feature and conjunction still better explained the dissimilarity of the stimulus reward probabilities as reflected in the difference between the weights of the informative feature and conjunction and the weight of object being larger than 0 (median±IQR = 0.43±0.31, two-sided sign-rank test; *P* < 10^−10^, *d* = 0.56, *N* = 50).

Altogether, results of lesioning suggest that the observed mixed feature- and conjunction-based learning strategies mainly rely on recurrent connections from inhibitory populations with plastic sensory input to excitatory populations with plastic sensory input. We next tested the importance of learning in these connections using alternative network architectures.

### Plasticity in recurrent connections is crucial for emergence of complex learning strategies

We trained three alternative architectures of RNNs to further confirm the importance of recurrent connections and reward-dependent plasticity in sensory and recurrent connections in our model. Specifically, we trained RNNs without plastic sensory input, RNNs without plastic recurrent connections, and feedforward neural networks (FFNNs) with only excitatory units and plastic sensory input.

We found that the RNNs without plastic sensory input were not capable of learning even non-structured stimulus-outcome associations during the training step. Moreover, although the RNNs without plastic recurrent connections and the FFNNs were able to perform the task and learn stimulus-outcome associations, their behavior was significantly different from the behavior of our participants. Specifically, the RNNs without plastic recurrent connections only exhibited an object-based learning strategy (**Supplementary Figure 10A**). In contrast, the explained variance of fit of FFNNs’ estimates over time was best fit by the F+C_1_ model (**Supplementary Figure 10B**), however, the time course of learning in this model was different from that exhibited by human participants. Specifically, unlike our experimental results, we did not find any significant difference between the time constant of increase in the weight of the informative feature and the time constant of increase in the weight of the informative conjunction in the FFNN (*τ_inf. feature_* : median±IQR = 37.04±29.09, *τ_inf. conjunction_* : median±IQR = 50.18±30.51, two-sided sign-rank test; *P* = 0.34, *d* = 0.47, *N* = 50; **Supplementary Figure 10C**). Together, these results demonstrate the importance of plasticity in recurrent connections between excitatory and inhibitory populations for the observed evolution of learning strategies.

## Discussion

Using a combination of experimental and modeling approaches, we investigated the emergence and adoption of value representations and learning strategies in more naturalistic settings. We show for the first time, that in high-dimensional environments, human participants estimate reward probabilities associated with each of many stimuli by learning and combining estimates of reward probabilities associated with the informative feature and the informative conjunction. As predicted, feature-based learning is much faster than conjunction-based learning and emerges earlier. Analyses of connectivity pattern and response of units in the trained RNNs illustrate that such mixed representations of reward value emerge over time as a distributed code that depends on distinct contributions of inhibitory and excitatory neurons. More specifically, learning about multidimensional stimuli results in contrasting connectivity patterns and representations of feature and object values in inhibitory and excitatory neurons, respectively. Through disinhibition, the recurrent connections allow gradual emergence of mixed (feature-based followed by conjunction-based) learning strategies in excitatory neurons. This emergence relies more strongly on connections from inhibitory to excitatory units because in the absence of these connections, object-based learning can quickly dominate. Our results thus, provide clear testable predictions about the emergence and neural mechanisms of naturalistic learning.

Our behavioral results support a previously proposed adaptability-precision tradeoff (APT) framework (Farashahi, Donahue, et al., 2017; Farashahi, Rowe, et al., 2017; Khorsand & Soltani, 2017) for understanding competition between different learning strategies. Moreover, they confirm our hypothesis that the complexity of learning strategies depends on the generalizability of the reward environment. In the absence of any instruction, human participants were able to detect the level of generalizability of the environment and learn the informative feature and conjunctions of non-informative features. This conjunction-based learning that follows learning about individual features allows the participants to improve accuracy in their learning without significantly compromising the speed of learning, thus improving the APT. This happens because feature-based, conjunction-based, and object-based learning allow updating of reward probabilities for 3 out of 9 individual features, 3 out of 27 possible conjunctions, and 1 out of 27 individual stimuli in each trial of the experiment, making conjunction-based learning the second fastest and mixed feature- and conjunction-based learning the second-most accurate strategy.

We found that the timescale at which conjunction-based learning emerged was an order of magnitude slower than that of feature-based learning. This result confirms the aforementioned notion about the rate of updates in different learning strategies and provides a critical test for finding the underlying neural architecture. Our results also indicate that humans can learn higher-order associations (conjunctions) when lower-order associations (non-informative features) are not useful. However, this only happens if the environment is stable enough (relative to the timescales of different learning strategies) such that there is sufficient time for slower representations to emerge before reward contingencies change. The findings on the timescale of different learning strategies have import implications for learning in naturalistic settings and for identifying their neural substrates (Spitmaan et al., 2020).

The categorization learning and stereotyping literature can provide an alternative but complementary interpretation of our behavioral results (Anderson, 1991; Ashby & Maddox, 2005; Gluck & Bower, 1988; Love et al., 2004). A task with generalizable reward schedule can be considered a rule-based reasoning task (i.e., a task in which optimal strategy can be described verbally), whereas a task with a non-generalizable reward schedule can be interpreted as an information integration task (i.e., a task in which information from two or more stimulus components should be integrated). The Competition between Verbal and Implicit Systems (COVIS) model of category learning assumes that rule-based category learning is mediated primarily by an explicit (i.e., hypothesis-testing) system, whereas information integration is dominated by an implicit (i.e., procedural-learning-based) system. According to COVIS, these two learning systems are implemented in different regions of the brain, but the more successful system gradually dominates (Ashby et al., 1998). In this framework, our results and the APT can be seen as a way of quantifying what factors can affect one learning strategy to dominate the other.

We found distributed representations of reward value in our proposed RNNs. Such representations have been the focus of many studies in memory (Rissman & Wagner, 2012), face- and object-encoding (O’toole et al., 2005; Pinsk et al., 2005; Tsao et al., 2003), and semantic knowledge (Small et al., 1995; Tyler & Moss, 2001), but have not been thoroughly examined in reward learning. Due to similarities between learning and categorization tasks, recurrent connections between striatum and prefrontal cortex have been suggested to play an important role in this process (Ashby et al., 1998; Radulescu et al., 2019). However, future experimental studies are required to investigate the emergence of distributed value representations in the reward learning system.

Our modeling results can also explain a few existing neural observations. First, we showed that recurrent inputs from certain inhibitory populations in the RNNs result in disinhibition of excitatory populations during the learning. Our RNNs structure does not make any specific assumption on where such disinhibition is originating from, and therefore it can be further extended to explain the effects of disinhibition in subcortical areas, such as the basolateral amygdala (Carlen et al., 2012; Letzkus et al., 2011, 2015; Wolff et al., 2014) or striatum (Lee et al., 2017; Owen et al., 2018), in associative learning. Second, our findings could explain a previously reported link between disruption in recurrent connections between excitatory and inhibitory neurons in basal ganglia and deficits in adoption of proper learning strategies in Parkinson’s Disease (Ashby et al., 2003; Ell et al., 2010; Price et al., 2009; Taverna et al., 2008). Finally, we observed a higher degree of similarity between activity of inhibitory recurrent units and the informative feature value, which is in line with recent findings on the prevalence of feature-specific error signal in narrow-spiking neurons (Oemisch et al., 2019). In addition, our results predict that error signals in neurons with plastic sensory input are skewed toward conjunction-based learning strategies.

Our experimental paradigm and computational model have a few limitations. First, because of the difficulty of the task, we used a fixed multi-dimensional reward environment. However, humans and animals are often required to learn reward values in changing environments. Future experiments and modeling are required to explore learning in dynamic multi-dimensional environments. Second, recent studies have suggested that reward can enhance processing of behaviorally relevant stimuli by influencing different low-level and high-level processes such as sensory representation, perceptual learning, and attention (Chen et al., 2015; Donahue & Lee, 2015; Goltstein et al., 2018; Khan et al., 2018; Oemisch et al., 2019; Poort et al., 2015; Ranganathan et al., 2018; Soltani et al., 2016; Spitmaan et al., 2019). Our RNNs only incorporate low-level synaptic plasticity, and thus, future studies are needed to understand the contribution of high-level processes (such as attention) in naturalistic value-based learning. Finally, we utilized a backpropagation algorithm to train the RNNs, which often is viewed as non-biological. Our aim was to offer a general framework to test hypotheses about learning in naturalistic environments. Nonetheless, recent studies have proposed dendritic segregation and local interneuron circuitry as a possible mechanism that can approximate this learning method (Guerguiev et al., 2017; Lillicrap et al., 2020; Roelfsema & Holtmaat, 2018; Sacramento et al., 2018; Whittington & Bogacz, 2019). Future progress in mapping the backpropagation method to learning in cortical structures would clarify the plausibility of this training approach.

Together, our study provides insight into both why and how more complex learning strategies emerge over time. On the one hand, we show that mixed learning strategies are adopted because they can provide an efficient intermediate learning strategy that is still much faster than learning about individual stimuli or options but is also more precise than feature-based learning alone. On the other hand, we provide clear testable predictions about neural mechanisms underlying naturalistic learning and thus, address both representations and functions of different neural types and connections.

## Acknowledgments

This work is supported by the National Science Foundation (CAREER Award BCS1943767 to A.S.). We would like thank Jane Xu for collecting some of the experimental data and Venice Nomof and Zohra Aslami for collecting data on an earlier version of the experiment.

## Methods

### Participants

Participants were recruited from the Dartmouth College student population. In total, 92 participants were recruited (66 females) and performed the experiment. We excluded participants whose performance was not significantly different from chance (0.5), an indication of not learning the task. To that end, we used a performance threshold of 0.55, equal to 0.5 plus 2 times s.e.m., based on the average of 400 trials after excluding the first 32 trials of the experiment. This resulted in exclusion of 25 participants from our dataset. We also analyzed the behavior of excluded participants to show that indeed they did not learn the task (**Supplementary Figure 4**). No participant had a history of neurological or psychiatric illness. Participants were compensated with a combination of money and “t-points,” which are extra-credit points for classes within the Department of Psychological and Brain Sciences at Dartmouth College. The base rate for compensation was $10/hour or 1 t-point/hour. Participants were then additionally rewarded based on their performance by up to $10/hour. All experimental procedures were approved by the Dartmouth College Institutional Review Board, and informed consent was obtained from all participants before participating in the experiment.

### Experimental paradigm

To study how appropriate representations of reward value and learning strategies are formed and evolve over time, we designed a novel probabilistic learning task in which participants learned about visual stimuli with three distinct features (shape, pattern, and color) by choosing between pairs of stimuli followed by reward feedback, and moreover, reported their learning about those stimuli in separate sets of trials (**Supplementary Figure 1**). More specifically, participants completed one session of 432 choice task (**Supplementary Figure 1A**) and five estimation bouts (after trials 86, 173, 259, 346, and 432 of the choice task; **Supplementary Figure 1C**). In each trial of the choice task, participants were presented with a pair of stimuli and were asked to choose the stimulus that they believed would provide the most reward. These two stimuli were drawn pseudo-randomly from a set of 27 stimuli, which were constructed using combinations of three distinct shapes, three distinct patterns, and three distinct colors. The two stimuli presented on each trial always differed in all three features. Selection of a given stimulus was rewarded (independently of the other presented stimulus) based on a reward schedule (set of reward probabilities) with a moderate level of generalizability. That is, reward probability associated with some but not all stimuli could be estimated by combining the reward probabilities associated with their features (see Eq. 1 below). More specifically, only one feature (shape or pattern) was informative about reward probability whereas the other two were not informative. Although the two non-informative features were on average not predictive of reward, specific combinations or conjunctions of these two features were partially informative of reward (**Supplementary Figure 1B**). Finally, during each estimation block, participants were asked to provide an estimate about the reward probabilities associated with individual stimuli (**Supplementary Figure 1C**).

### Data analysis

Unless otherwise mentioned, the statistical comparisons were performed using Wilcoxon signed-rank test in order to test the hypothesis of zero median for one sample or the difference between paired samples. The reported effect sizes are Cohen’s *d*-values. All behavioral analyses and model fitting were done using MATLAB 2018a (MathWorks, Inc., Natick, MA). RNN simulations were implemented using the tensor flow package in Python 3.7 environment.

### Reward schedule

The reward schedule (matrix) was constructed to allow testing the adoption of different learning strategies. Assume that stimuli or objects (*0*) have *m* features (e.g., color, pattern, and shape), each of which can take *n* different instances (e.g., yellow, solid, and triangles), indicated as *F_ij_* for the feature instance *j* of feature *i*, where *i* = {1,…,*m*} and *j* = {1,…,*n*}. A feature-based learner can use the average reward probability for each feature instance to estimate reward probability associated with each stimulus in two steps. First, the average reward probability for a given feature instance (e.g., color yellow) can be computed by averaging the reward probability of all stimuli that contain that feature instance (e.g., all yellow stimuli); 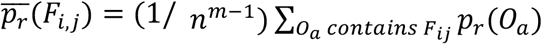 or by multiplying the likelihood ratios of all stimuli that contain that feature instance: 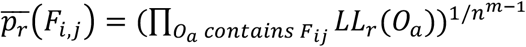. Second, reward probability for a stimulus 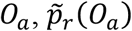, can be estimated by combining the reward probability of features of that stimulus using the Bayes theorem:

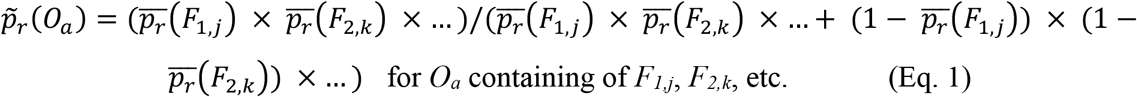

These estimated reward probabilities constitute the *estimated reward matrix* based on features. The rank order of probabilities in the estimated reward matrix, which determines preference between stimuli, is similar to that of the fully generalizable reward matrix, whereas the exact probabilities may differ slightly.

Similarly, the mixed feature- and conjunction-based learner can combine the reward probability for one or more of feature instances, *F_ij_* (where *i*=*I* ⊆ {1,…,*m*} and *j* = {1,…,*n*}), and the conjunctions of the other remaining features (e.g., solid triangle, indicated as *C_l,k_* for the conjunction instance *k, k* = {1,…,*n*^*m*|*I*|^}) of possible conjunctions of features *l* (where *l_i_* ={1,…,*m*}-I) to estimate reward probabilities of stimuli in three steps. First, the average reward probability associated with one or more feature instances can be calculated as above. Second, the average reward probability associated with one or more conjunctions of remaining features can be computed by averaging the reward probabilities of all stimuli that contain that conjunction instance (e.g., all solid triangle stimuli); 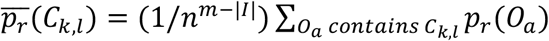, or by multiplying the likelihood ratios of all stimuli that contain that conjunction instance; 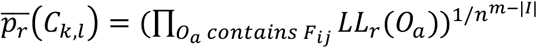. Finally, reward probability for a stimulus 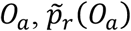, can be estimated by combining the reward probabilities of features and conjunctions using the Bayes theorem:

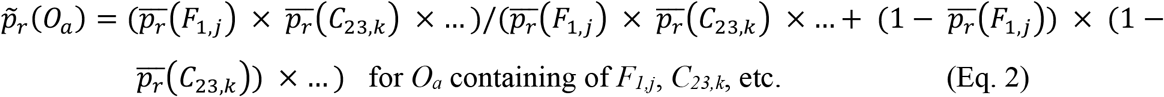

### Generalizability index

To define generalizability indices, we used the correlation between the actual reward probability of stimuli and the estimated reward probability of stimuli based on their individual features, or between actual reward probability of stimuli and estimated reward probability of stimuli based on the mixture of the informative feature and the conjunctions of the two non-informative features. On the basis of this definition, the generalizability index can take on any values between −1 and 1.

### Using estimates of reward probabilities to assess learning strategies

We utilized the estimates participants provided during estimation sessions in order to examine how they determined the reward probabilities of stimuli. Specifically, we used GLMs to predict participants’ estimates of reward probabilities as a function of the following variables: actual reward probabilities assigned to each stimulus (object-based term); reward probability estimates based on the combination of the reward probabilities associated with individual features (feature-based term; Eq.1); reward probability estimates based on the combination of the reward probability of the informative feature and the conjunctions of the other two non-informative features (mixed feature- and conjunction-based term; Eq. 2); and a constant. The constant (bias) term in this GLM captures participants’ overall bias in reporting reward probabilities.

### Fitting the time course of regression weights for predicting participants’ estimates

To quantify the time course of learning using participants’ estimates, we fit the regression weights for the informative feature and conjunction using an exponential function:

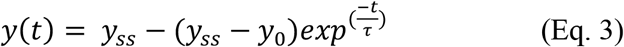

where *y*_0_ and *y_ss_* are the initial and steady-state values of weights, *τ* is the time constant for approaching steady state, and *t* represents the trial number in a session.

### Testing behavioral predictions of learning models

To directly assess the effects of learning strategy on choice behavior, we defined a few quantities we refer to as “differential response” separately for the informative and non-informative features and conjunctions of non-informative features. Differential response for individual features is defined as the difference between two conditional probabilities: 1) conditional probability of selecting stimuli that contain only one of the two features (informative or non-informative) of the stimulus selected and rewarded in the previous trial when these stimuli were paired with a stimulus that did not share any feature with the previously rewarded stimulus; and 2) a similar conditional probability when the previous trial was not rewarded (see **Supplementary Figure 5**). Similarly, we defined differential response for conjunctions of non-informative features by considering stimuli that contain the conjunction of non-informative features of the stimulus selected in the previous trial.

### Model fitting

To capture participants’ learning and choice behavior on a trial-by-trial basis, we used 24 different reinforcement learning (RL) models based on feature-based, mixed feature- and conjunction-based, and object-based strategies. These models were fit to experimental data during the choice task by minimizing the negative log likelihood of the predicted choice probability given different model parameters using the “fminsearch” function in MATLAB (Mathworks). We computed three measures of goodness-of-fit in order to determine the best model: average negative log likelihood, Akaike information criterion (AIC), and Bayesian information criterion (BIC). In addition, to compare the ability of different models in fitting choice behavior over time, we used AIC and BIC per trial (Farashahi et al., 2018, 2020), denoted as AIC_p_ and BIC_p_:

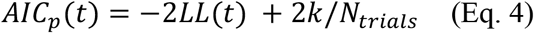

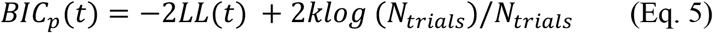

where *k*indicates the number of parameters in a given model, *t* represents the trial number, *LL*(*t*) is the log likelihood in trial *t*, and *N_trials_* is the number of trials in the experiment. By dividing the penalty terms in AIC and BIC by the number of trials, we ensure that the sum of AICp(*t*) and BICp(*t*) over all trials is equal to AIC and BIC, respectively. The smaller values for these measures indicate a better fit of choice behavior.

### Object-based models

In this group of models, the reward probability associated with each stimulus is directly estimated from reward feedback in each trial using a standard RL model. For example, in the uncoupled object-based RL, only the reward probability associated with the chosen stimulus is updated in each trial. This update is done via separate learning rates for rewarded or unrewarded trials using the following equations, respectively:

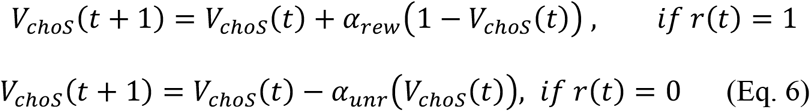

where *t* represents the trial number, *V_choO_* is the estimated reward probability associated with the chosen stimulus, *r*(*t*) is the trial outcome (1 for rewarded, 0 for unrewarded), and *α_rew_* and *α_unr_* are the learning rates for rewarded and unrewarded trials. The reward probability associated with the unchosen stimulus is not updated in this model.

In the coupled object-based RL, the reward probabilities associated with both stimuli presented in a given trial are updated, but in opposite directions. That is, while reward probability associated with the chosen stimulus is updated based on Eq. 6, the value of unchosen stimulus is updated based on the following equation:

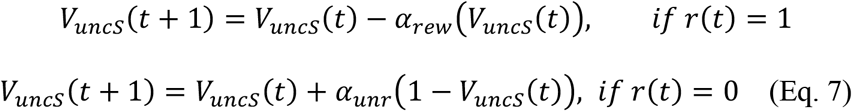

where *t* represents the trial number and *V_uncS_* is the estimated reward probability associated with the unchosen stimulus.

The estimated reward probability values are then used to compute the probability of selecting between the two stimuli in a given trial (*S1* and *S2*) based on a logistic function:

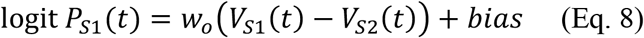

where *P_S1_* is the probability of choosing stimulus 1, *V_S1_* and *V_S2_* are the estimated reward probability associated with stimuli 1 and 2, respectively, *bias* measures a response bias toward the left or right option to capture the participant’s location bias, and *w_o_* determines the influence of reward probability associated with objects on choice.

### Feature-based models

In this group of models, the reward probability associated with each stimulus is computed by combining the reward probability associated with features of that stimulus, which are estimated from reward feedback using a standard RL model. The update rules for the feature-based RL models are identical to the object-based ones, except that the reward probability associated with the chosen (unchosen) stimulus is replaced by the reward probability associated with features of the chosen (unchosen) stimulus.

Similar to object-based RL models, the probability of choosing a stimulus is determined based on the logistic function of the difference between the estimated reward probability associated with the two presented stimuli in a given trial:

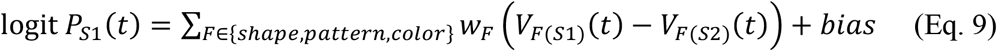

where *V*_*F*(*S*1)_ and *V*_*F*(*S*2)_ are the reward probabilities associated with a feature of stimuli 1 and 2, respectively, *bias* measures a response bias toward the left or right option to capture the participant’s location bias, and three values of *w_F_* determine the influence of reward probability associated with each feature on choice.

### Mixed feature- and conjunction-based models

In this group of models (referred to as the F+C models for abbreviation), the reward probability associated with each stimulus is computed by combining the reward probabilities associated with one feature and the conjunction of the other two features of that stimulus, all of which are estimated from reward feedback using a standard RL model. In these models, we assume that subjects identify one feature as the likely informative feature early in the experiment and subsequently learn about the conjunctions of the other two features. The update rules for the mixed feature- and conjunction-based RL models are identical to the previous models, except that the reward probability associated with the chosen (unchosen) stimulus is replaced by the reward probability associated with a feature or the conjunction of the other two features of the chosen (unchosen) stimulus. Additionally, we considered separate learning rates for rewarded and unrewarded trials (*α_rew_*, *α_unr_*) when updating reward probabilities associated with the feature and conjunction. This setting results in three different mixed feature- and conjunction-based models depending on which feature and conjunction of other two features are being learned. In the F+C_1_ model, we assume that the informative feature and the conjunction of the other two non-informative features are being learned; for example, values of shapes and conjunctions of colors and patterns are learned (**Supplementary Figure 1**). In the F+C_2_ and F+C_3_ models, one of the two non-informative features is learned along with the conjunction of the other non-informative feature and the informative feature. In the example reward schedule shown in Supplementary Figure 1, reward probabilities associated with colors and conjunctions of shapes and patterns are learned in F+C_2_, whereas reward probabilities associated with patterns and conjunctions of shapes and colors are learned in F+C_3_.

Finally, the probability of choosing a stimulus is determined based on the logistic function of the difference between the total estimated reward probabilities associated with two presented stimuli in a given trial:

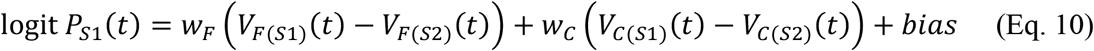

where *V*_*F*(*S*1)_ and *V*_*F*(*S*2)_ are reward probabilities associated with a feature of stimuli 1 and 2, respectively, *V*_*C*(*S*1)_ and *V*_*C*(*S*2)_ are the reward probabilities associated with the conjunction of the other two features of stimuli 1 and 2, respectively, *bias* measures a response bias toward the left or right option to capture the participant’s location bias, and *w_F_* and *w_C_* determine the influence of the reward probability associated with features and conjunctions on choice, respectively.

### Mixed feature- and object-based models

In this group of models (referred to as F+O models for abbreviation), the reward probability associated with each stimulus is computed by combining the reward probabilities associated with one feature and the reward probability associated with stimulus directly estimated from reward feedback in each trial using a standard RL model. In these models, we assume that subjects identify one feature as the likely informative feature early in the experiment and combine this learning with what they learn about individual stimuli later in the experiment. The update rules for the mixed feature- and object-based RL models are identical to the previous models, except that the reward probability associated with the chosen (unchosen) stimulus is replaced by the reward probability associated with a feature or the chosen (unchosen) stimulus itself. Additionally, we considered separate learning rates for rewarded and unrewarded trials (*α_rew_*, *α_unr_*) when updating reward probabilities associated with the feature and stimulus. This setting results in three different mixed feature- and object-based models depending on which feature is learned along with objects. In the F_1_+O model, we assume that the informative feature and the objects are being learned; for example, values of shapes and objects are learned (**Supplementary Figure 1**). In the F_2_+O and F_3_+O models, the non-informative features are learned along with the value of the objects. In the example reward schedule shown in Supplementary Figure 1, reward probabilities associated with colors and objects are learned in F_2_+O whereas reward probabilities associated with patterns and objects are learned in F_3_+O.

Finally, the probability of choosing a stimulus is determined based on the logistic function of the difference between the total estimated reward probabilities associated with two presented stimuli in a given trial:

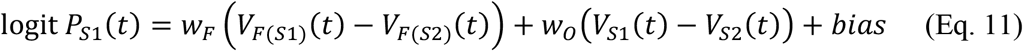

where *V*_*F*(*S*1_) and *V*_*F*(*S*2)_ are reward probabilities associated with a feature of stimuli 1 and 2, respectively, *V*_*S*1_ and *V*_*S*2_ are the reward probabilities associated with the stimuli 1 and 2, respectively, *bias* measures a response bias toward the left or right option to capture the participant’s location bias, and *w_F_* and *w_O_* determine the influence of the reward probability associated with features and objects on choice, respectively.

### RL models with decay

Additionally, we investigated the effect of “forgetting” reward probabilities associated with the unchosen objects or feature(s) (for the uncoupled models) by introducing a decay in estimated probabilities that has been shown to capture some aspects of learning, especially in multi-dimensional tasks (Farashahi, Rowe, et al., 2017; Niv et al., 2015). More specifically, reward probabilities associated with unchosen stimuli or feature(s) decay to 0.5 with a rate of *d*, as follows:

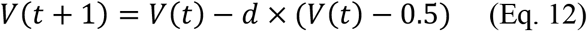

where *t* represents the trial number and *V* is the estimated reward probability associated with a feature, a stimulus, or a conjunction of two features.

### Recurrent neural network

To understand computations and neural mechanisms underlying the emergence of different learning strategies, we used recent methods for training recurrent neural networks (Mante et al., 2013; Masse et al., 2019; Rajan et al., 2016; Song et al., 2016, 2017; Wang et al., 2018; Yang et al., 2019) to construct biophysically-plausible recurrent networks of point neurons endowed with plausible reward-dependent Hebbian learning rules (Gerstner et al., 2014; Loewenstein & Seung, 2006; Pfeiffer et al., 2010). Specifically, we used stochastic gradient descent (SGD) to train recurrent neural networks (RNNs) consisting of excitatory and inhibitory units both to learn input-output associations for estimating reward probabilities and to perform our learning task in two separate steps.

In the first (training) step, we trained RNNs to learn the temporal dynamics of input-output associations and a set of reward probabilities randomly associated with 27 stimuli used in our task. This was done to enable the network to learn a universal solution for learning reward probabilities in environments with different levels of generalizability. We used the stochastic gradient descent (SGD) method to train the network to learn reward probabilities using a three-factor, reward-dependent Hebbian learning rule for weights from sensory units to recurrent units and between recurrent units. Reward probabilities were drawn from a uniformly random distribution between 0 and 1 for each training session that consisted of 270 trials. The RNNs were trained until their performance (MSE of the estimates at the end of the session) with reward probabilities similar to our experimental paradigm matched the average performance of our participants. In the second step (simulation of the experiment), we stopped SGD and simulated the behavior of the trained RNNs in a session with reward probabilities similar to our experimental paradigm and compared the behavior of the RNNs with that of our participants. In this step, only plastic connections were modulated after each reward feedback.

The overall structure of the simulated task was similar to our experimental paradigm with a few exceptions. Specifically, only one stimulus was shown in each trial and the network had to learn the reward probability associated with that stimulus. Moreover, in each trial, the network received visual information about the presented stimulus (i.e., distinct sensory inputs representing features, conjunction of features, and object-identity) as a tonic input and had to learn reward probability associated with that stimulus using reward feedback provided in each trial. We did not require our network to demonstrate working memory to be able to better isolate necessary structure for multi-dimensional reward learning. Using this approach, we avoided complexity related to decisionmaking processes and mainly focused on learning aspects of our experimental paradigm.

### Network structure

RNNs received a set of *N*_in_ = 63 time-varying inputs (*u_t_*) and were trained to produce an output *z_t_*, where inputs encoded stimuli-relevant sensory information and output represented the estimate of the network for reward probability. Network input consisted of nodes representing all features, conjunctions of features, and the object-identity of the stimuli. Network contained *N* = 120 units that learn the association between input and the output (**Figure 3**). The activity of recurrent units followed the continuous dynamical equation as below (capital letters refer to matrixes and small letters refer to vectors and scalars):

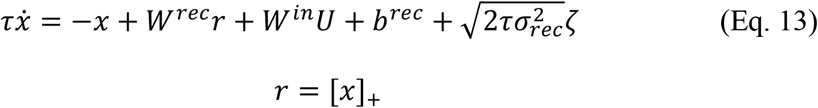

where *τ* is the time constant of the activity in the recurrent units, *W^in^* is an *N*_in_×*N* matrix of connection weights from the sensory units (*u*) to recurrent units, *W^rec^* is an *N*×*N* matrix of recurrent connection weights, *b^rec^* is a bias term, *ζ* is Gaussian white noise processes with zero mean and unit variance, *σ_rec_* is the strength of the intrinsic noise to the network, and []_+_ is the rectifying linear function (ReLu). An output unit *z* reads out nonlinearly from the network as below:

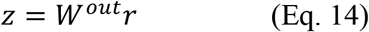

where *W^out^* is an *1×N* matrix of connection weights from the recurrent units to the output. To solve the aforementioned dynamical system, we used the first-order Euler approximation with a time-discretization step Δ*t* to arrive at the following equations:

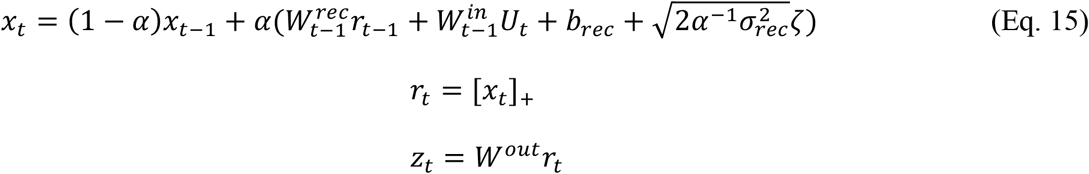

where 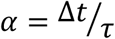 and *t* = [0, T] is the time elapsed within a single trial. The network received noisy sensory input from 63 populations of sensory encoding units (*U_t_*) as follows:

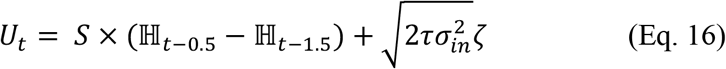

where ℍ is the Heaviside function allowing stimuli onset at 0.5*s* and stimuli offset at 1.5*s*, *S* is a vector with 1s corresponding to feature, conjunctions and the object-identity of the presented stimulus, and *σ_in_* is the strength of the input noise. In our simulations, we set Δ*t* = 0.02*s*, *τ* = 0.1*s*, *σ_rec_* = 0.01, and *σ_in_* = 0.01.

Critically, we designed our network to obey biological constraints observed in mammalian cortex. Specifically, we constrained units to have purely excitatory or inhibitory effects and hypothesized that excitatory units (*Exc*) outnumber inhibitory units (*Inh*) by a ratio of 4 to 1. In addition, we assumed that all inputs and outputs of the network were long-range excitatory inputs from an upstream circuit, and therefore all elements of 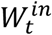 and *W^out^* were forced to be non-negative. To impose excitatory or inhibitory effects on recurrent connections, we implemented 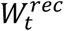 as the product between a matrix for which all entries were trained and forced to be non-negative (*W*^*rec*+^) and a fixed diagonal matrix with 1s corresponding to excitatory units and −1s corresponding to inhibitory units on its main diagonal (Masse et al., 2019; Song et al., 2016).

To enable the network to learn a universal structure of our task, we embedded three-factor reward-dependent Hebbian learning in selected weights of the RNNs (Gerstner et al., 2014). Specifically, we hypothesized that a combination of pre- and post-synaptic activity (*F*(*pre_i_, post_j_*)) and the modulator signal (*D_t_*) results in an update of weights as follows:

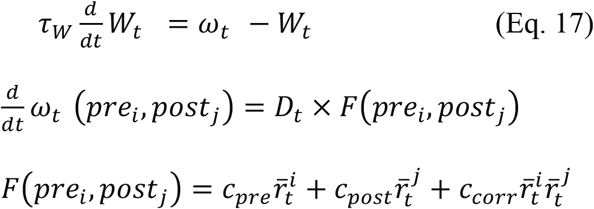

where *c_pre_, c_post_, c_corr_* are the pre-, post-synaptic, and the correlation terms, 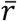 represents the firing rate low-pass filtered with time constant *τ_r_* = 0.1s, and *τ_w_ =* 0.1s is the time constant of the weights’ update. The modulator signal *D_t_* was defined as follows:

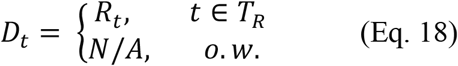

where *T_R_* = [1.75*s*, 2*s*] represents the time when reward feedback is present, and *R_t_* denotes the binary reward (+1 when reward was present and −1 when reward was absent).

To systematically examine the contribution of different recurrent units to RNNs’ behavior, we assigned each unit in the inhibitory or excitatory population to one of four types of populations: (a) *Exc_rr_* and *Inh_rr_* corresponding to populations with no plastic sensory or recurrent connections (rigid weights indicated by subscript *r*); (b) *Exc_fr_* and *Inh_fr_* corresponding to populations with plastic sensory input only (flexible weights indicated by subscript *f*); (c) *Exc_rf_* and *Inh_rf_* corresponding to populations with plastic recurrent connections only; and (d) *Exc_ff_* and *Inh_ff_* corresponding to populations with plastic sensory input and plastic recurrent connections. Finally, we simulated three alternative versions of our model. We trained RNNs without plastic sensory input, RNNs without plastic recurrent connections, and feedforward neural network (FFNNs) with only excitatory units with plastic sensory input.

### RNN training procedure

In the first “training” step, we used the Adam version of SGD (Kingma & Ba, 2014) to train our RNNs. The default set of parameters were used for the training, where the learning rate was 0.001 and the decay rate for the first- and second-moment estimates were 0.9 and 0.999, respectively. The objective function 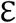 to be minimized was computed by time-averaging the squared errors between the network output *z_t_* and the target output 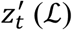 in addition to using the *L*_2_-norm regularization term (*R*) for encouraging sparse weights/activation patterns. The objective function was calculated as follows:

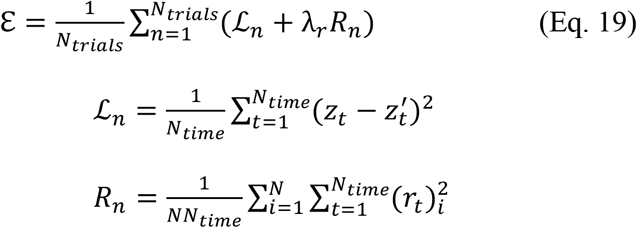

where *λ_r_* = 0.01 determines the effect of the regularization term *R_n_*. During training, we adjusted 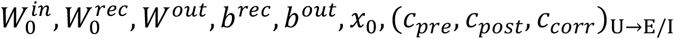, and (*c_pre_*, *c_post_*, *c_corr_*)_E/I→E/I_ parameters, where *W*_0_ is the initial connection weights and *x*_0_ refers to the initial neuronal activity at time *t* = 0 in each trial. The network was trained with randomly generated reward probabilities between 0 and 1 for each session where target output 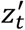 was defined as the reward probability of the stimuli during the choice period (*T_ch_* = [1*s*, 1.5*s*]) and zero activity before the stimuli onset ([0*s*, 0.5*s*]). Each training session consisted of *N_trials_* = 270 trials of *T* = 2*s*, equivalent to a session in our experimental paradigm. Networks’ performance, defined as MSE of estimates at the end of a session with reward probabilities equal to our experimental paradigm, was used as the training cutoff. Training was continued until each network achieved a final estimation error similar to that observed in participants’ last reported estimates.

### Simulating the experiment with the trained RNNs

In the second step, we stopped SGD and simulated the behavior of the trained RNNs in a session with reward probabilities equal to our experimental paradigm. Specifically, during these simulations 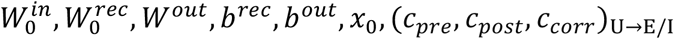 and (*c_pre_*, *c_post_*, *c_corr_*)_E/I→E/I_ parameters were fixed at the values found in the training step, while plastic connections were modulated after each reward feedback according to Eqs. 17–18.

### Analysis of response in recurrent units

To identify the representations of different strategies in our network model, we examined the dissimilarity in the response of recurrent units to all stimuli and its relationship to dissimilarity of reward value of different stimuli based on different learning strategies (Hunt et al., 2018; Kriegeskorte et al., 2008). The “response dissimilarity matrix” was computed as the distance between the activity of recurrent units in each population during the choice period. The “reward probability dissimilarity matrix” was calculated as the distance between reward probability estimates based on each of the three models (informative-feature, conjunction of noninformative features, and object-based models) for all the stimuli. We then used GLMs to estimate the normalized weights of the reward probability dissimilarity matrix for predicting the response dissimilarity matrix. We obtained similar results when measuring dissimilarity with Euclidian distance, correlation, and cosine, but only report those based on the Euclidian distance.

### Measuring reward modulations of connection weights

To measure the effects of reward-dependent plasticity, we examined changes in plastic sensory input weights during the simulation of our task. Specifically, we used GLM to fit weights from a specific sensory unit type (e.g., yellow-color encoding units) to specific recurrent units (e.g., *Exc_fr_*) using the reward value estimates associated with those sensory units (value of color yellow). Value estimates were calculated using a simple reinforcement learning agent (learning rates (*α_rew_*, *α_unr_*) = 0.05). Additionally, we used GLMs to estimate changes in plastic recurrent connection weights as a function of time (trial number).

## Supplemental material

**Supplementary Figure 1.**
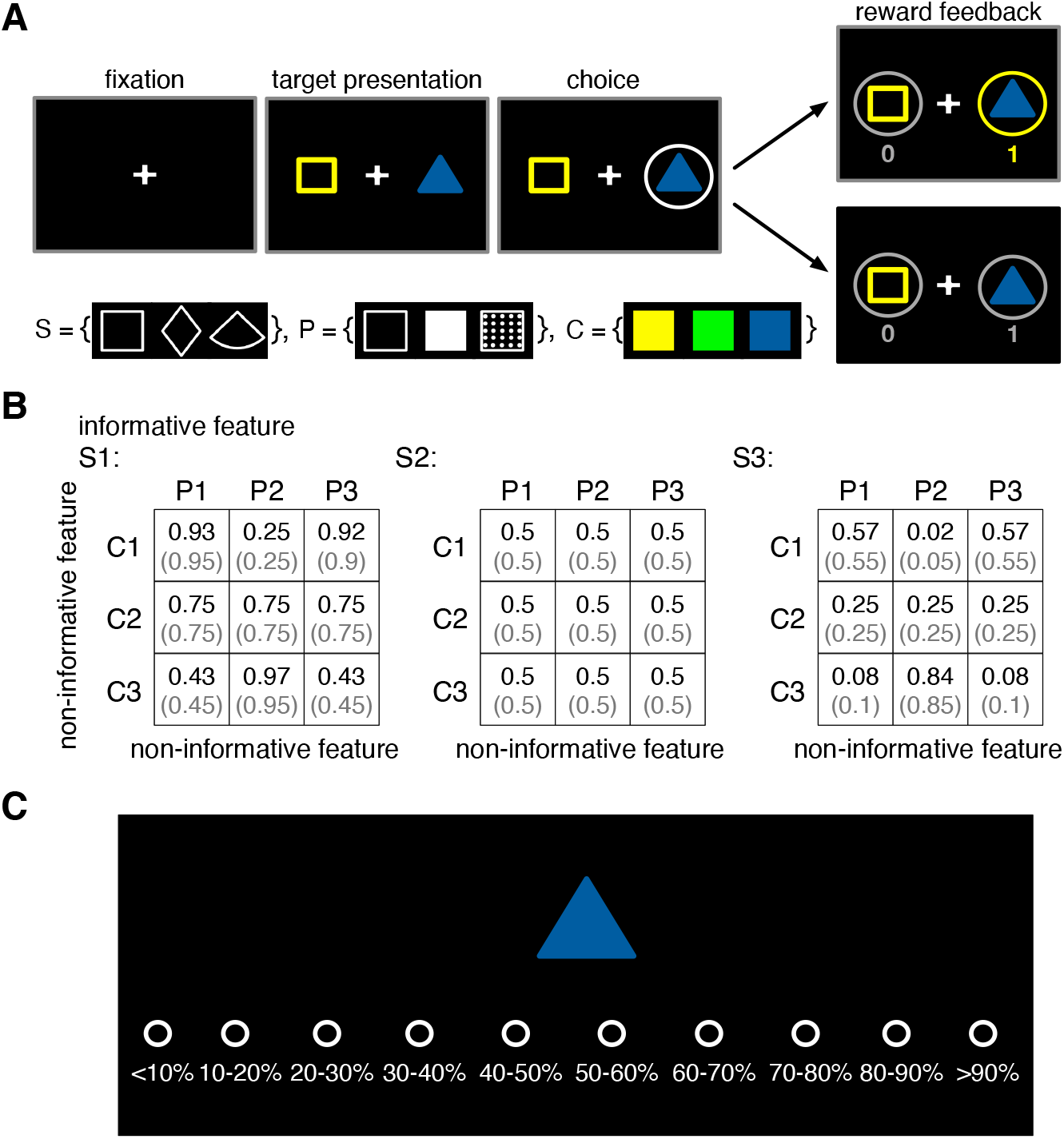
Experimental procedure. **(A)** Timeline of a trial during the multi-dimensional probabilistic learning task. In each trial, the participant chose between two stimuli (colored patterned shapes) and was provided with reward feedback (reward or no reward) for the chosen stimulus. The inset at the bottom shows the set of all visual features used in our experimental paradigm (S, shape; P, pattern; C, color). **(B)** Example of reward probabilities assigned to 27 possible stimuli. Stimuli were defined by combinations of three features, each with three instances. Reward probabilities were non-generalizable, such that reward probabilities assigned to all stimuli could not be determined by combining the reward probabilities associated with their features or conjunctions of their features. Numbers in parentheses demonstrate the actual probability values used in the experiment due to limited resolution for reward assignment. For the example schedule, the shape was on average informative about reward (average probability of reward on three shapes were equal to 0.3, 0.5, and 0.7). However, although pattern and color alone were not informative (average probability of reward for these features was equal to 0.5, 0.5, and 0.5), the conjunction of two non-informative features was on average informative about the reward (average probability of reward for conjunctions was equal to {0.63, 0.27, 0.63; 0.5, 0.5, 0.5; 0.37, 0.74, 0.37}). Each participant was randomly assigned to a condition where the informative feature was either pattern or shape. (**C**) A sample estimation trial. On each estimation trial, the participant estimated the probability of reward on an individual stimulus by pressing one of ten keys (A, S, D, F, G, H, J, K, L, and ;) on the keyboard.

**Supplementary Figure 2.**
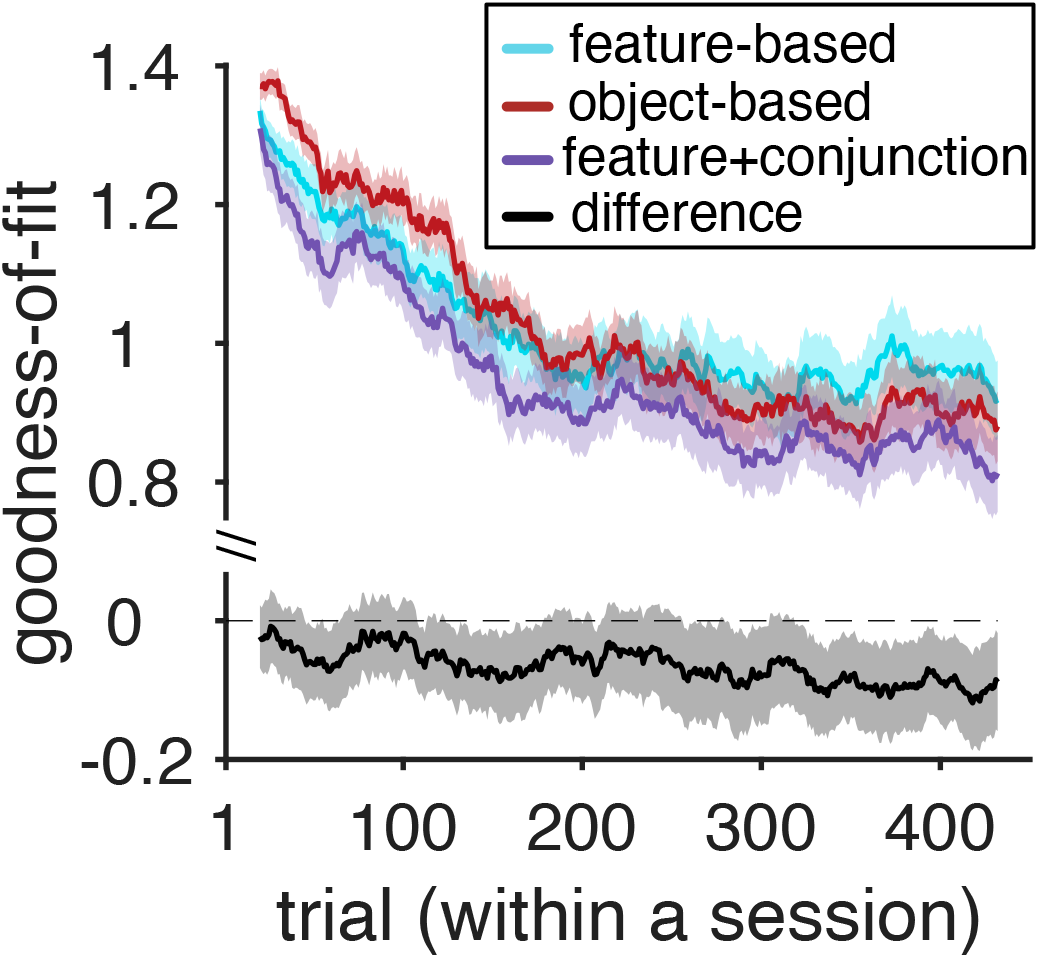
Plotted is the goodness-of-fit based on the average BIC per trial, BICp, for the feature-based model, object-based model, and the best mixed feature- and conjunction-based (F+C_1_) model. The smaller value corresponds to a better fit. The black curve shows the difference between the goodness-of-fit for the F+C_1_ model and the feature-based model.

**Supplementary Figure 3.**
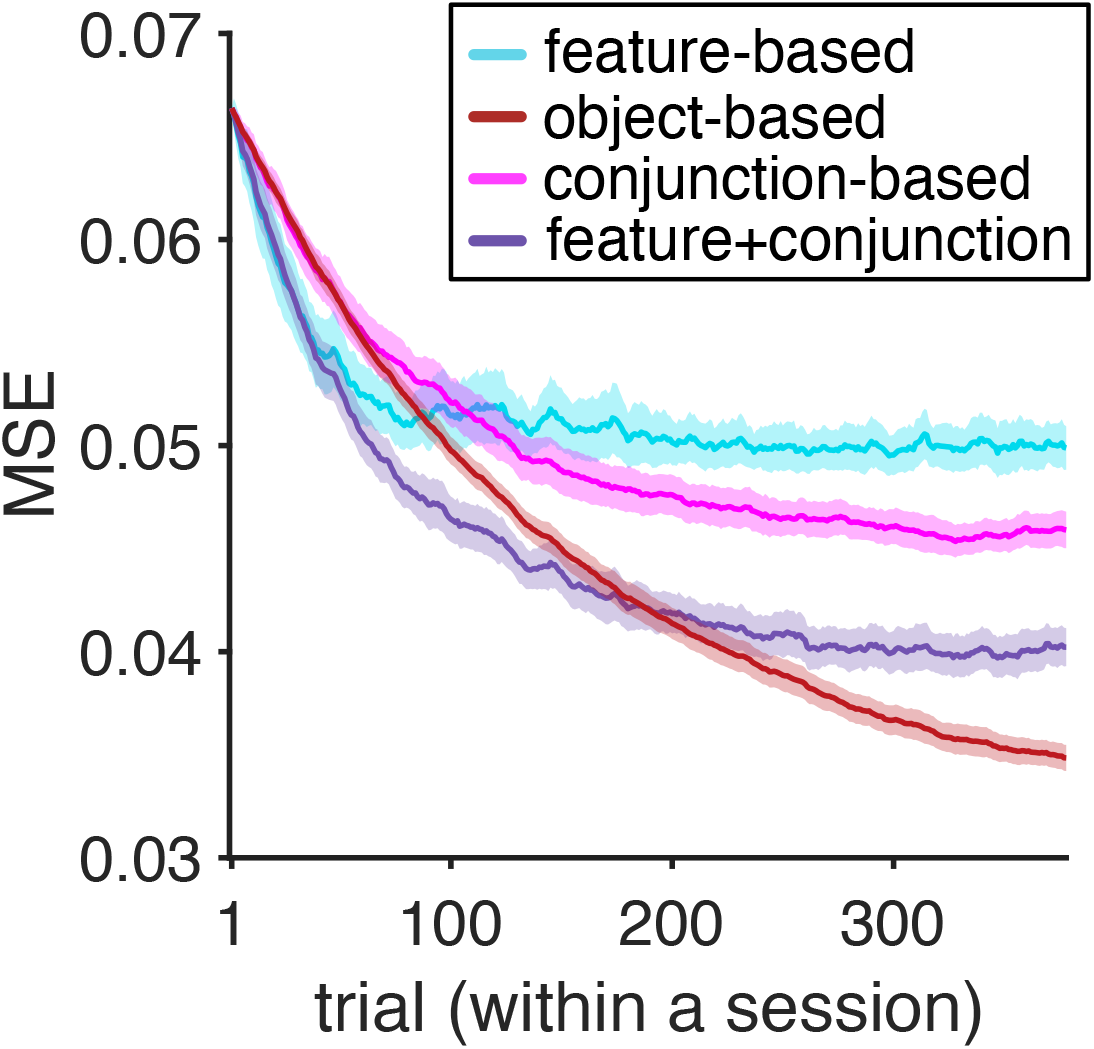
Average squared error (MSE) in the estimation of reward probabilities using a reinforcement learning model with decay (the learning rates (*α_rew_*, *α_unr_*) = 0.05 and the decay rate *d* = 0.005) and based on different learning strategies. A feature-based learner and a mixed feature- and conjunction-based learner (F+C_1_) have a lower MSE at the beginning of the experiment, whereas an object-based learner provides more accurate estimates later in the experiment. The shaded areas indicate s.e.m.

**Supplementary Figure 4.**
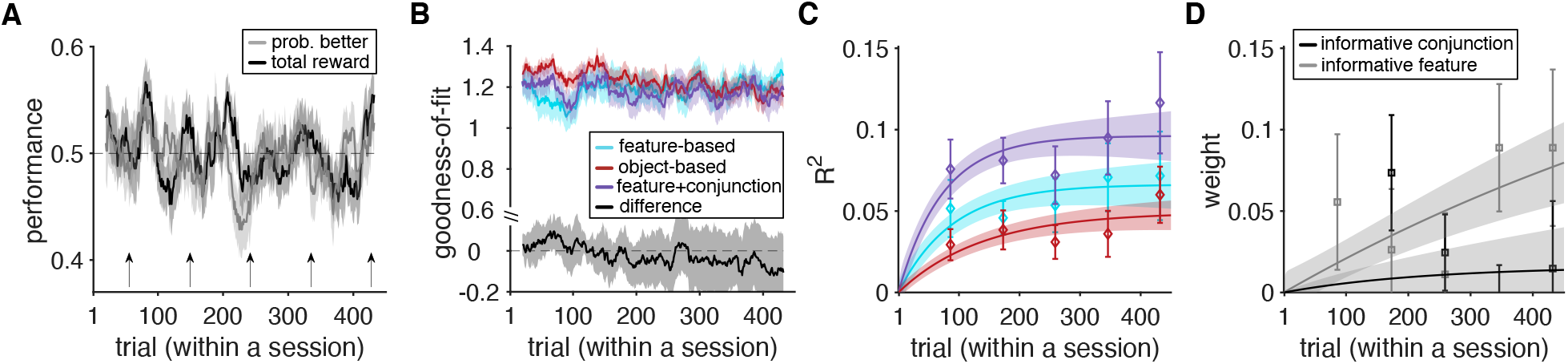
Analyses of choice behavior and estimation of excluded participants (*N* = 25). (**A**) Time course of performance and learning during the experiment. Plotted are the total harvested reward and probability of selecting the stimulus with higher probability of reward (better option) in a given trial within a session of the experiment. The running average over time is computed using a moving box with the length of 20 trials. Shaded areas indicate s.e.m., and the dashed line shows chance performance. Overall, these participants failed to learn reward probabilities associated with the options. (**B**) Plotted is the goodness-of-fit based on the average AIC per trial, AICp, for the feature-based model, object-based model, and the best mixed feature- and conjunction-based (F+C_1_) model. Similar results hold using average BIC per trial, BICp (data not shown). The smaller value corresponds to a better fit. The black curve shows the difference between the goodness-of-fit for the F+C_1_ and feature-based models. Throughout most of the experiment, the F+C_1_ model provided a better fit for choice behavior of excluded participants, similarly to that for the participants included in the study. (**C**) The plot shows the time course of explained variance (*R^2^*) in participants’ estimates based on different GLMs. Color conventions are the same as in panel B with cyan, red, and purple curves representing *R^2^* based on feature-based, object-based, and the F+C_1_ models, respectively. The solid line is the average of fitted exponential function to each participant’s data, and shaded areas indicate s.e.m. of the fit. (**D**) Time course of adopted learning strategies measured by fitting participants’ estimates of reward probabilities. Plotted is the weight of the informative feature and informative conjunction in the F+C_1_ model. Error bars represent s.e.m. The solid line is the average of fitted exponential function to each participant’s data, and shaded areas indicate s.e.m. of the fit. Overall, we found no evidence that the excluded participants adopted a strategy qualitatively different from the one used by the rest of the participants.

**Supplementary Figure 5.**
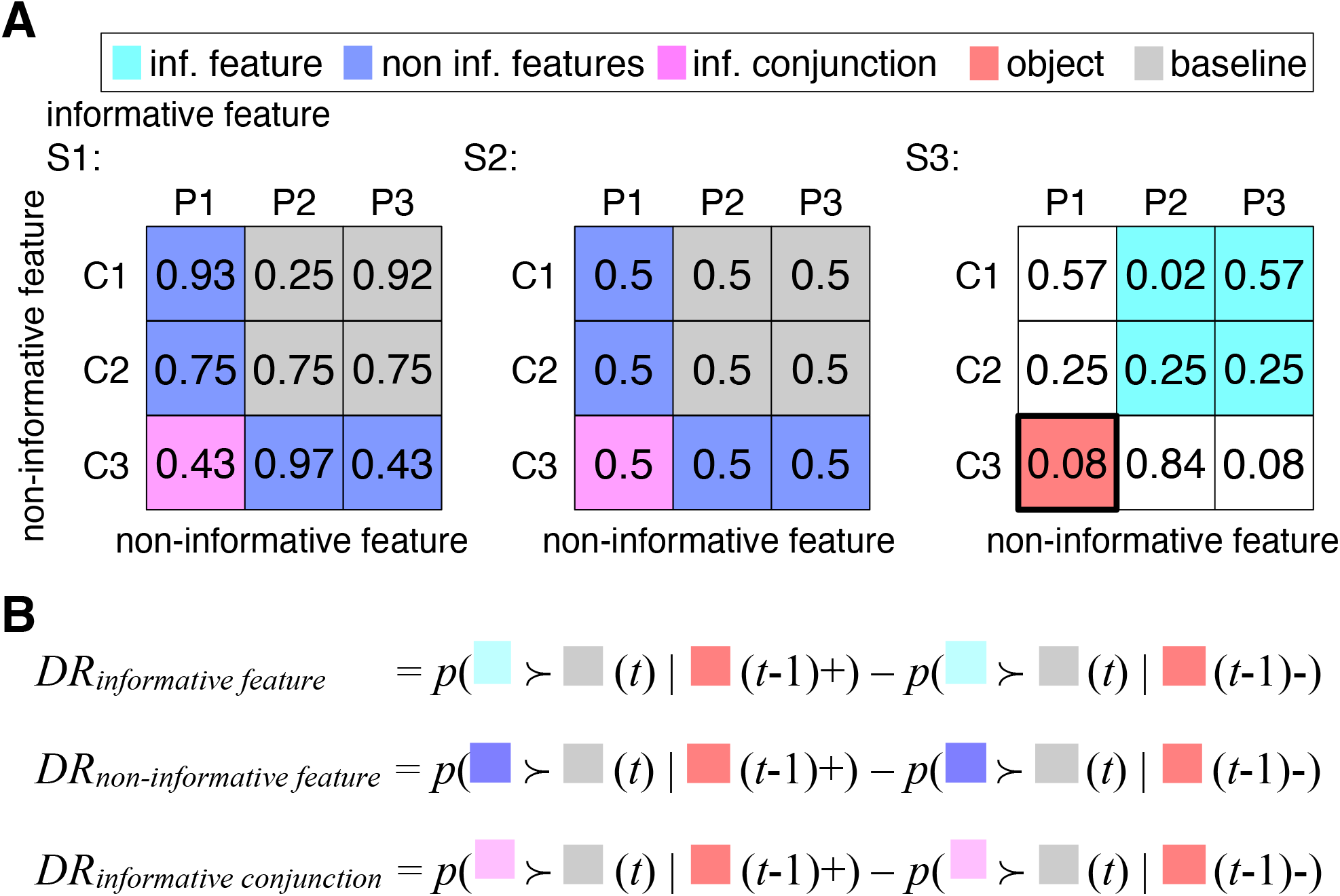
Differential response to reward feedback as a proxy for learning strategy. (**A**) Illustration of stimuli with common features and conjunctions. For an example of chosen stimulus in trial *t* ({*S3, P1, C3*} highlighted in red), we show stimuli that share only the informative feature, the non-informative features, or the conjunction of non-informative features in cyan, blue, and magenta, respectively. Stimuli depicted in gray do not share any common features or conjunction of features with the previously chosen stimulus. Reward probabilities are similar to those in Supplementary Figure 1. (**B**) Rules used to calculate differential responses. The probability of choosing *X* when it is presented together with *Y* in trial *t* given that choosing *Z* was rewarded and unrewarded in the previous trial is denoted as *p(X≻Y(t)|Z(t-1)+)* and *p(X≻Y(t)|Z(t-1)-)*, respectively. Differential response for the informative or non-informative features is defined as the probability of selecting stimuli that contains only the informative or non-informative features of the stimulus selected and rewarded in the previous trial minus the same probability when the previous trial was not rewarded. Similarly, differential response for conjunction of non-informative features is calculated for stimuli that contain the conjunction of non-informative features of the stimulus selected in the previous trial. All these probabilities were calculated for trials where the chosen stimulus on trial *(t-1)* was paired with a stimulus that did not share any features or conjunction of features with the previously chosen stimulus.

**Supplementary Figure 6.**
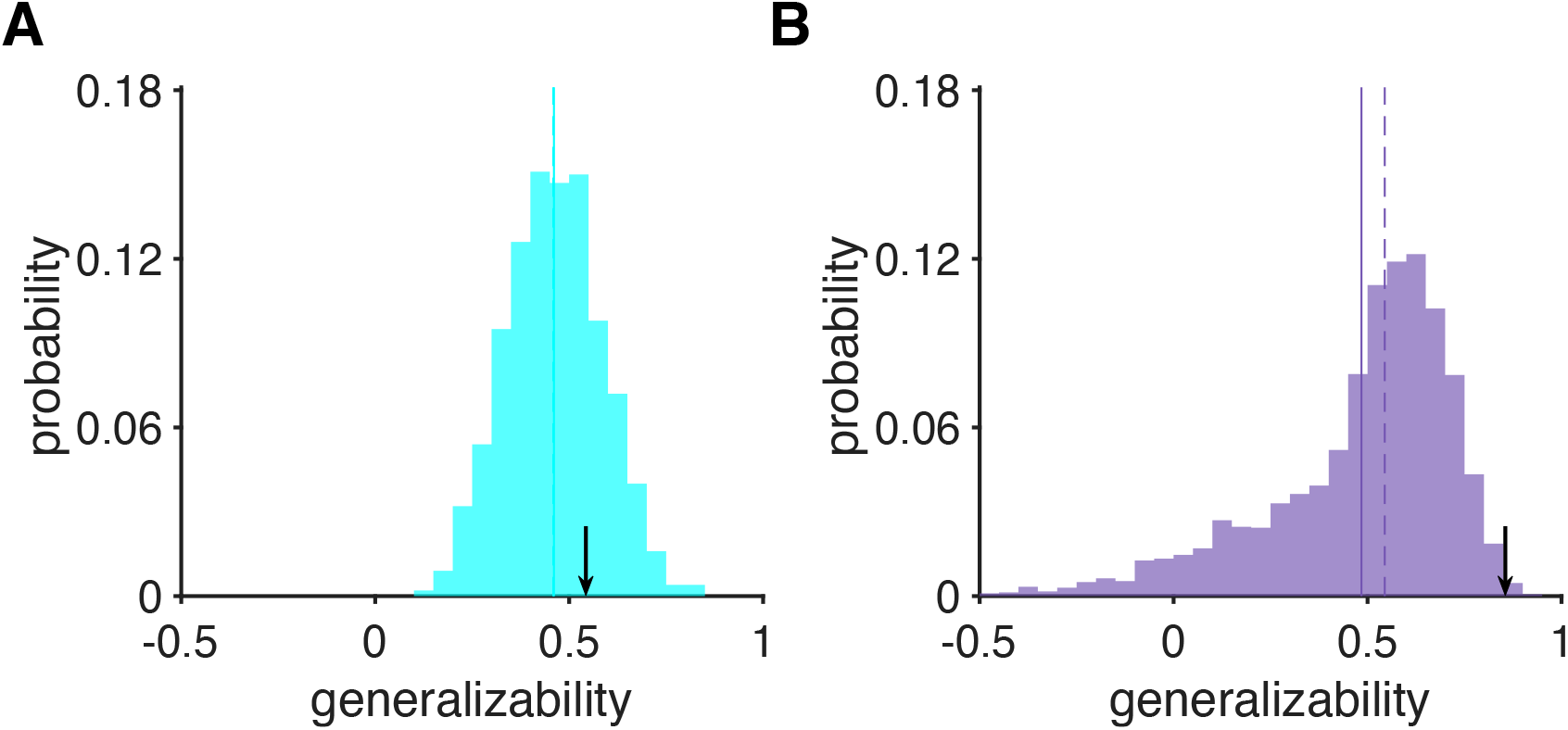
Distribution of generalizability indices in the environments used for training our RNNs. The plots show the distribution of generalizability indices calculated for the estimated reward probabilities associated with different stimuli based on their features (A) and the mixture of individual features and conjunctions (B). The dashed and solid lines show the mean and median of the distributions, respectively. The arrows show the generalizability values associated with the reward schedule used in our task.

**Supplementary Figure 7.**
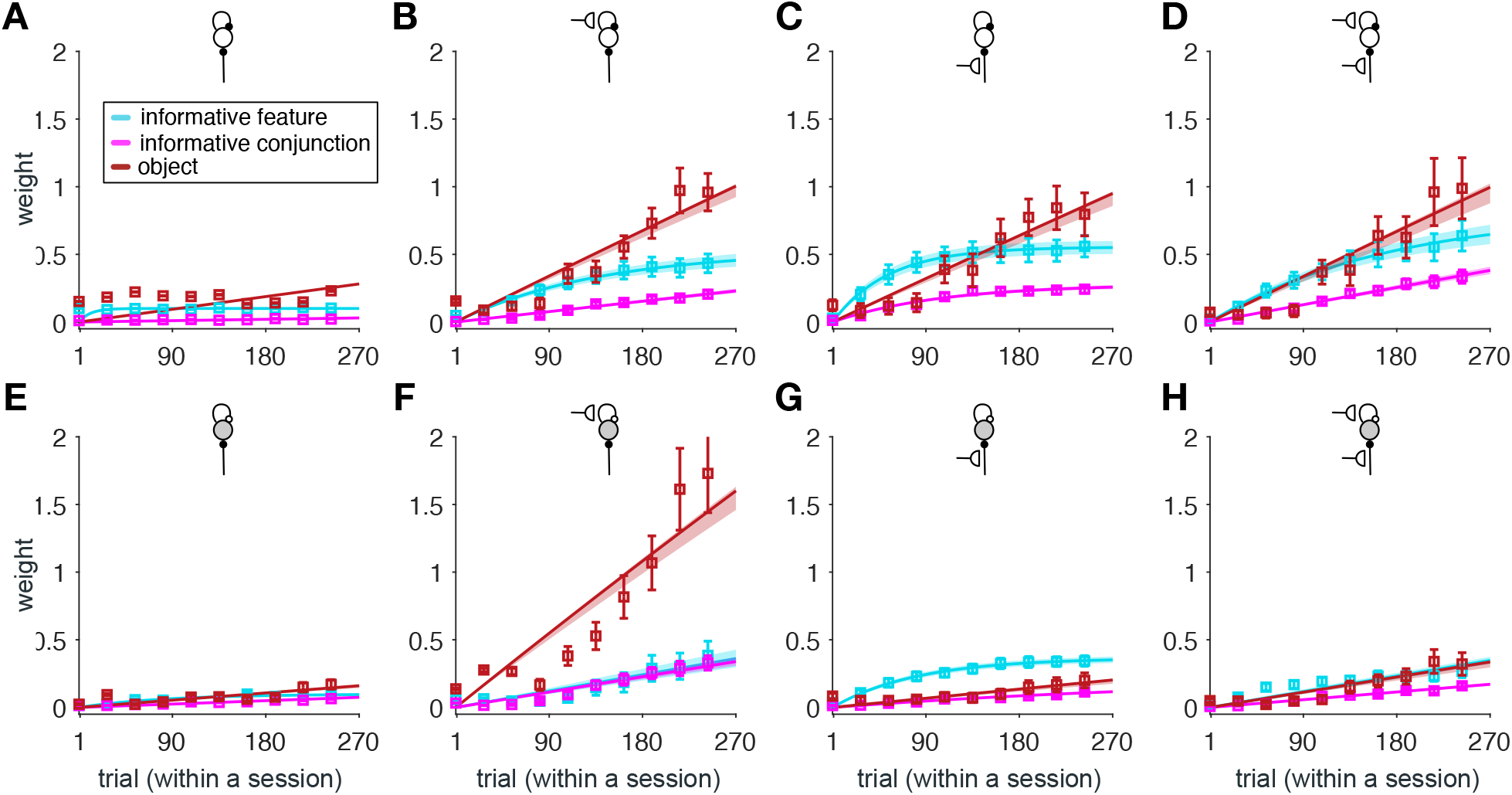
Similarity of response in different recurrent populations to reward probabilities based on different learning strategies for the RNNs in which recurrent connections from inhibitory populations with plastic sensory inputs to excitatory populations with plastic sensory inputs are lesioned. (**A–D**) Plotted are the estimated weights for predicting the response dissimilarity matrix of different types of recurrent populations (indicated by the inset diagrams explained in Figure 3B) using the dissimilarity of reward probabilities based on the informative feature, informative conjunction, and object. Error bars represent s.e.m. The solid line is the average of fitted exponential functions to RNNs’ data, and the shaded areas indicate s.e.m. of the fit. (**E–H**) Same as A–D but for inhibitory recurrent populations.

**Supplementary Figure 8.**
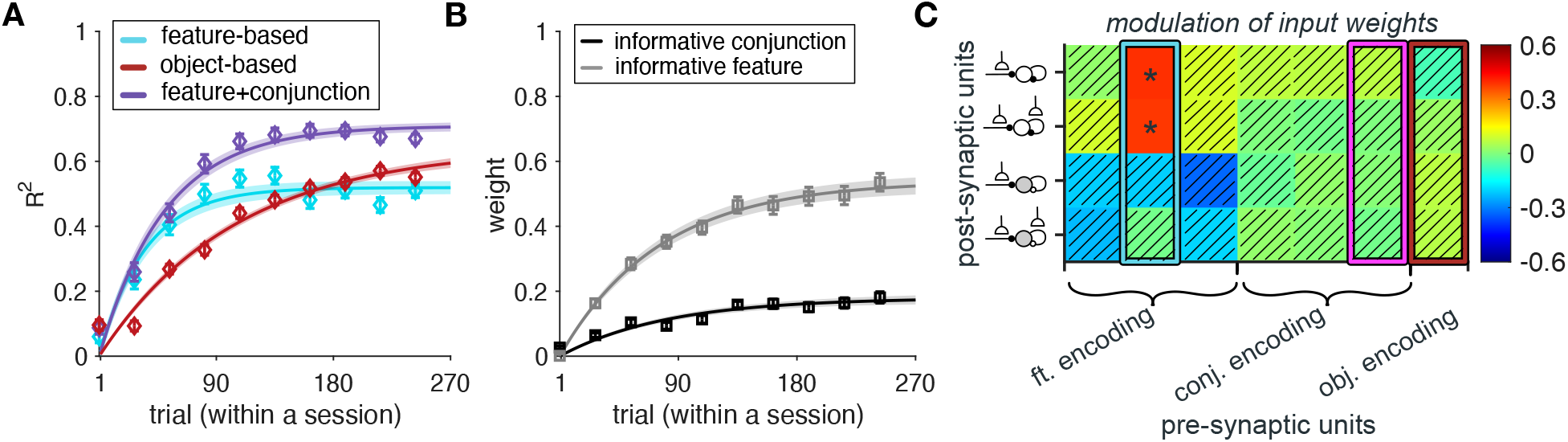
Results of lesioning connections from the excitatory populations with plastic sensory input (*Exc_fr_* and *Exc_ff_*) to the inhibitory populations with plastic sensory input (*Inh_fr_* and *Inh_ff_*) in the RNNs. (**A**) The plot shows the time course of explained variance (*R^2^*) in RNNs’ estimates based on different models. Error bars represent s.e.m. The solid line is the average of exponential fits to RNNs’ data, and the shaded areas indicate s.e.m. of the fit. (**B**) Time course of adopted learning strategies measured by fitting the RNNs’ output. Plotted is the weight of the informative feature and the informative conjunction in the F+C_1_ model (*τ_inf.feature_* = 76.34, *τ_inf.conjunction_* = 91.08). Error bars represent s.e.m. The solid line is the average of exponential fits to RNNs’ data, and the shaded areas indicate s.e.m. of the fit. (**C**) Plotted is the average rate of value-dependent changes in the connection weights from feature-encoding, conjunction-encoding, and object-identity encoding units to recurrent units with plastic sensory input during the simulation of our task. Asterisks indicate significant rates of change (*P*<0.05), and hatched squares indicate connections with rates of change that were not significantly different from zero (*P*>0.05). Highlighted rectangles in cyan, magenta, and red indicate the values for input from sensory units encoding the informative feature, the informative conjunction, and object-identity, respectively.

**Supplementary Figure 9.**
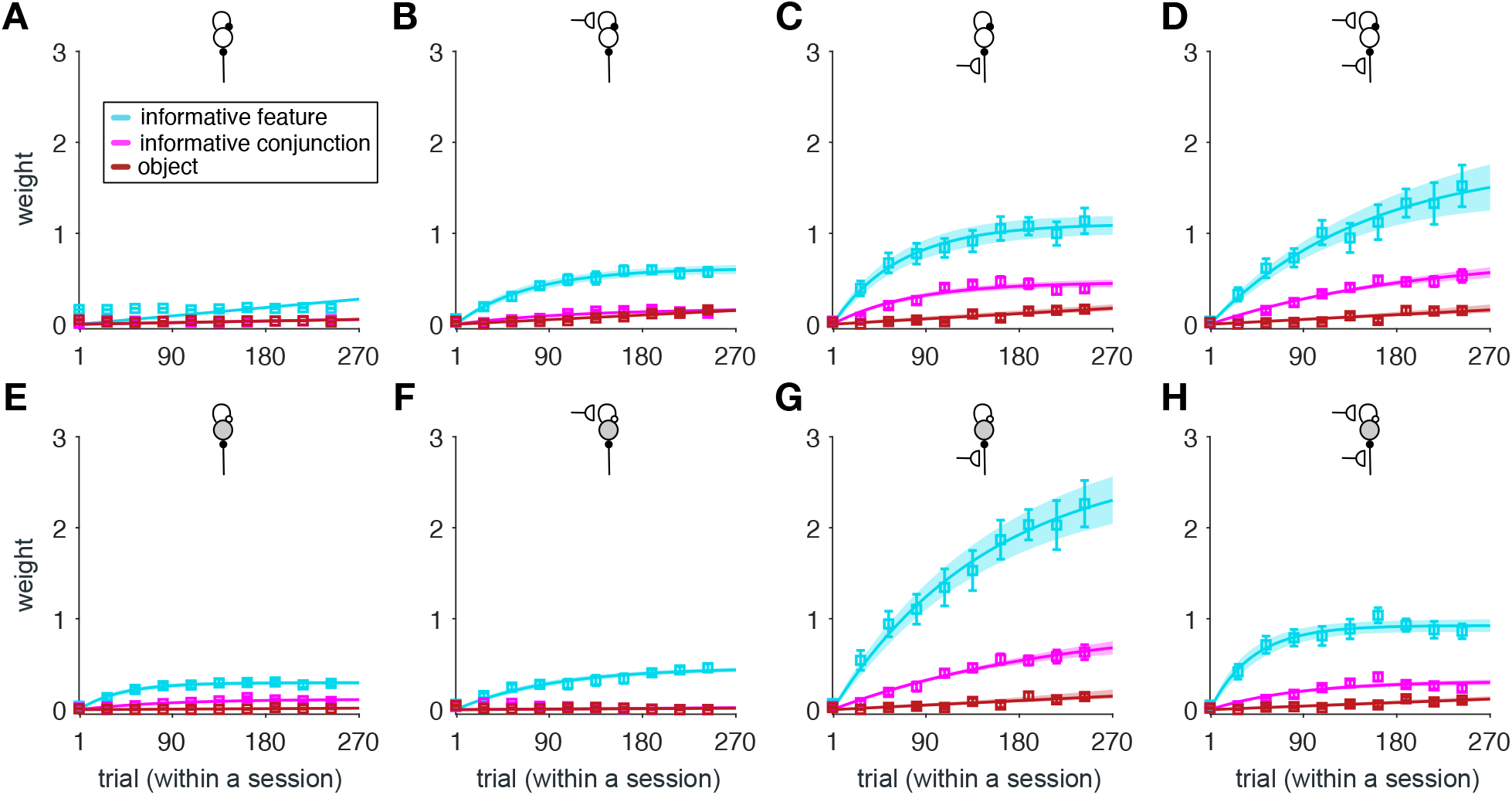
Similarity of response in different recurrent populations to reward probabilities based on different learning strategies for the RNNs in which recurrent connections from excitatory populations with plastic sensory input to inhibitory populations with plastic sensory input are lesioned. (**A–D**) Plotted are the estimated weights for predicting the response dissimilarity matrix of different types of recurrent populations (indicated by the inset diagrams explained in Figure 3B) using the dissimilarity of reward probabilities based on the informative feature, informative conjunction, and object. Error bars represent s.e.m. The solid line is the average of fitted exponential functions to RNNs’ data, and the shaded areas indicate s.e.m. of the fit. (**E–H**) Same as A-D but for inhibitory recurrent populations.

**Supplementary Figure 10.**
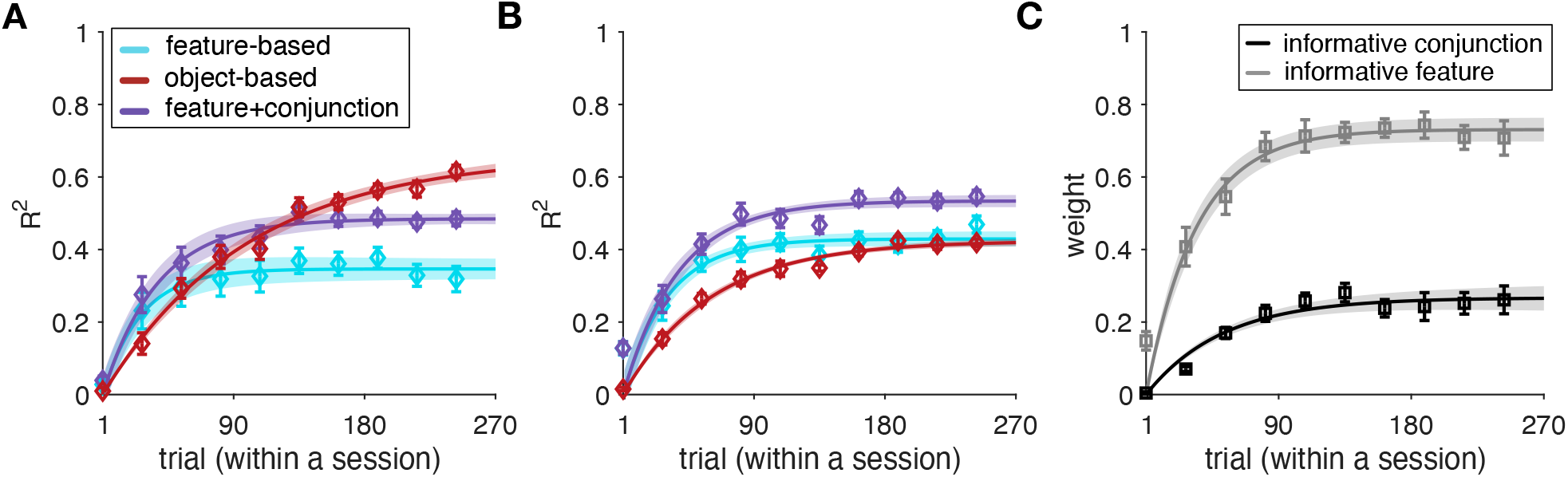
RNNs without reward-dependent plasticity in recurrent connections and feedforward neural networks (FFNNs) fail to replicate experimental results. (**A–B**) The plots show the time course of explained variance (*R^2^*) in RNNs’ estimates based on different models for the RNNs without reward-dependent plasticity in recurrent connections (A) and FFNNs (B). Error bars represent s.e.m. The solid line is the average of exponential fits to RNNs’ data, and the shaded areas indicate s.e.m. of the fit. (**C**) Time course of adopted learning strategies measured by fitting the RNNs’ output. Plotted is the weight of informative feature and the informative conjunction in the F+C_1_ model for FFNNs (*τ_inf.feature_* = 37.04, *τ_inf.conjunction_* = 50.18). Error bars represent s.e.m. The solid line is the average of exponential fits to RNNs’ data, and the shaded areas indicate s.e.m. of the fit.

**Supplementary Table 1.**
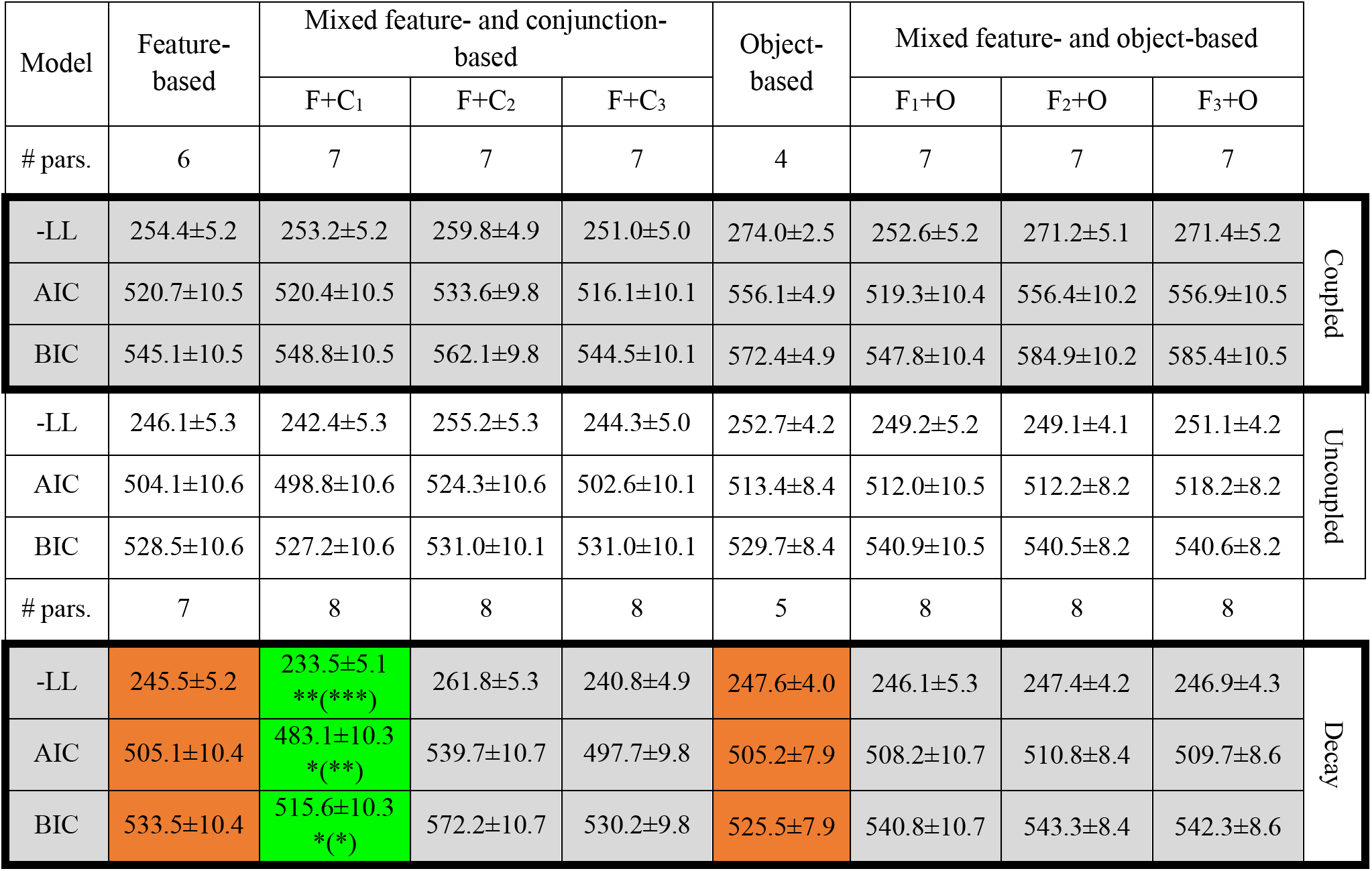
Comparison of the goodness-of-fit measures in the experiment. Reported are the goodness-of-fit measures, negative log likelihood (-LL), Akaike information criterion (AIC), and Bayesian information criterion (BIC) averaged over all participants (mean±s.e.m.) for different groups of models. The mixed model providing the best fit (F+C_1_) and its object-based and feature-based counterparts are highlighted in green and orange, respectively. The symbols next to goodness-of-fit for F+C_1_ indicate comparison with the feature-based and object-based (shown in parenthesis) models using a two-sided, signrank test. The significance level of the test is coded as: 0.01 < *P* < 0.05 (*), 0.001 < *P* < 0.01 (**), and *P* < 0.001 (***).

## References

Anderson, J. R. (1991). The adaptive nature of human categorization. Psychological Review, 98(3), 409.

Ashby, F. G., Alfonso-Reese, L. A., Waldron, E. M., & others. (1998). A neuropsychological theory of multiple systems in category learning. Psychological Review, 105(3), 442.

Ashby, F. G., & Maddox, W. T. (2005). Human category learning. Annu Rev Psychol, 56, 149–178.

Ashby, F. G., Noble, S., Filoteo, J. V., Waldron, E. M., & Ell, S. W. (2003). Category learning deficits in Parkinson’s disease. Neuropsychology, 17(1), 115.

Barlow, H. B. (1972). Single units and sensation: A neuron doctrine for perceptual psychology? Perception, 1(4), 371–394.

Braun, D. A., Mehring, C., & Wolpert, D. M. (2010). Structure learning in action. Behavioural Brain Research, 206(2), 157–165.

Carlen, M., Meletis, K., Siegle, J., Cardin, J., Futai, K., Vierling-Claassen, D., Ruehlmann, C., Jones, S. R., Deisseroth, K., Sheng, M., & others. (2012). A critical role for NMDA receptors in parvalbumin interneurons for gamma rhythm induction and behavior. Molecular Psychiatry, 17(5), 537–548.

Carnevale, F., de Lafuente, V., Romo, R., Barak, O., & Parga, N. (2015). Dynamic control of response criterion in premotor cortex during perceptual detection under temporal uncertainty. Neuron, 86(4), 1067–1077.

Chen, J. L., Margolis, D. J., Stankov, A., Sumanovski, L. T., Schneider, B. L., & Helmchen, F. (2015). Pathway-specific reorganization of projection neurons in somatosensory cortex during learning. Nature Neuroscience, 18(8), 1101.

Dayan, P., & Berridge, K. C. (2014). Model-based and model-free Pavlovian reward learning: Revaluation, revision, and revelation. Cognitive, Affective, & Behavioral Neuroscience, 14(2), 473–492.

Decharms, R. C., & Zador, A. (2000). Neural representation and the cortical code. Annual Review of Neuroscience, 23(1), 613–647.

Donahue, C. H., & Lee, D. (2015). Dynamic routing of task-relevant signals for decision making in dorsolateral prefrontal cortex. Nature Neuroscience, 18(2), 295–301.

Eliasmith, C., Stewart, T. C., Choo, X., Bekolay, T., DeWolf, T., Tang, Y., & Rasmussen, D. (2012). A large-scale model of the functioning brain. Science, 338(6111), 1202–1205.

Ell, S. W., Weinstein, A., & Ivry, R. B. (2010). Rule-based categorization deficits in focal basal ganglia lesion and Parkinson’s disease patients. Neuropsychologia, 48(10), 2974–2986.

Farashahi, S., Donahue, C. H., Khorsand, P., Seo, H., Lee, D., & Soltani, A. (2017). Metaplasticity as a neural substrate for adaptive learning and choice under uncertainty. Neuron, 94(2), 401–414.

Farashahi, S., Rowe, K., Aslami, Z., Gobbini, M. I., & Soltani, A. (2018). Influence of learning strategy on response time during complex value-based learning and choice. PloS One, 13(5), e0197263.

Farashahi, S., Rowe, K., Aslami, Z., Lee, D., & Soltani, A. (2017). Feature-based learning improves adaptability without compromising precision. Nature Communications, 8, 1768.

Farashahi, S., Xu, J., Wu, S.-W., & Soltani, A. (2020). Learning arbitrary stimulus-reward associations for naturalistic stimuli involves transition from learning about features to learning about objects. Cognition, 205, 104425.

Gershman, S. J., & Niv, Y. (2010). Learning latent structure: Carving nature at its joints. Current Opinion in Neurobiology, 20(2), 251–256.

Gerstner, W., Kistler, W. M., Naud, R., & Paninski, L. (2014). Neuronal dynamics: From single neurons to networks and models of cognition. Cambridge University Press.

Gluck, M. A., & Bower, G. H. (1988). From conditioning to category learning: An adaptive network model. Journal of Experimental Psychology: General, 117(3), 227–247.

Goltstein, P. M., Meijer, G. T., & Pennartz, C. M. (2018). Conditioning sharpens the spatial representation of rewarded stimuli in mouse primary visual cortex. Elife, 7, e37683.

Goudar, V., & Buonomano, D. V. (2018). Encoding sensory and motor patterns as time-invariant trajectories in recurrent neural networks. Elife, 7, e31134.

Guerguiev, J., Lillicrap, T. P., & Richards, B. A. (2017). Towards deep learning with segregated dendrites. ELife, 6, e22901.

Hinton, G. E. (1986). Learning distributed representations of concepts. Proceedings of the Eighth Annual Conference of the Cognitive Science Society, 1, 12.

Hinton, G. E., McClelland, J., & Rumelhart, D. (1986). Distributed representations. In: Parallel distributed processing: Explorations in the microstructure of cognition, vol. 2, Psychological and biological models.

Hunt, L. T., Malalasekera, W. N., de Berker, A. O., Miranda, B., Farmer, S. F., Behrens, T. E., & Kennerley, S. W. (2018). Triple dissociation of attention and decision computations across prefrontal cortex. Nature Neuroscience, 21(10), 1471–1481.

Khan, A. G., Poort, J., Chadwick, A., Blot, A., Sahani, M., Mrsic-Flogel, T. D., & Hofer, S. B. (2018). Distinct learning-induced changes in stimulus selectivity and interactions of GABAergic interneuron classes in visual cortex. Nature Neuroscience, 21(6), 851.

Khorsand, P., & Soltani, A. (2017). Optimal structure of metaplasticity for adaptive learning. PLoS Computational Biology, 13(6), e1005630.

Kingma, D. P., & Ba, J. (2014). Adam: A method for stochastic optimization. ArXiv Preprint ArXiv:1412.6980.

Kriegeskorte, N., Mur, M., & Bandettini, P. A. (2008). Representational similarity analysis-connecting the branches of systems neuroscience. Frontiers in Systems Neuroscience, 2, 4.

Lake, B. M., Salakhutdinov, R., & Tenenbaum, J. B. (2015). Human-level concept learning through probabilistic program induction. Science, 350(6266), 1332–1338.

Lake, B. M., Ullman, T. D., Tenenbaum, J. B., & Gershman, S. J. (2017). Building machines that learn and think like people. Behavioral and Brain Sciences, 40.

Lee, K., Holley, S. M., Shobe, J. L., Chong, N. C., Cepeda, C., Levine, M. S., & Masmanidis, S. C. (2017). Parvalbumin interneurons modulate striatal output and enhance performance during associative learning. Neuron, 93(6), 1451–1463.

Leong, Y. C., Radulescu, A., Daniel, R., DeWoskin, V., & Niv, Y. (2017). Dynamic interaction between reinforcement learning and attention in multidimensional environments. Neuron, 93(2), 451–463.

Letzkus, J. J., Wolff, S. B., & Lüthi, A. (2015). Disinhibition, a circuit mechanism for associative learning and memory. Neuron, 88(2), 264–276.

Letzkus, J. J., Wolff, S. B., Meyer, E. M., Tovote, P., Courtin, J., Herry, C., & Lüthi, A. (2011). A disinhibitory microcircuit for associative fear learning in the auditory cortex. Nature, 480(7377), 331–335.

Lillicrap, T. P., Santoro, A., Marris, L., Akerman, C. J., & Hinton, G. (2020). Backpropagation and the brain. Nature Reviews Neuroscience, 1–12.

Loewenstein, Y., & Seung, H. S. (2006). Operant matching is a generic outcome of synaptic plasticity based on the covariance between reward and neural activity. Proceedings of the National Academy of Sciences, 103(41), 15224–15229.

Love, B. C., Medin, D. L., & Gureckis, T. M. (2004). SUSTAIN: a network model of category learning. Psychological Review, 111(2), 309.

Mante, V., Sussillo, D., Shenoy, K. V., & Newsome, W. T. (2013). Context-dependent computation by recurrent dynamics in prefrontal cortex. Nature, 503(7474), 78–84.

Masse, N. Y., Yang, G. R., Song, H. F., Wang, X.-J., & Freedman, D. J. (2019). Circuit mechanisms for the maintenance and manipulation of information in working memory. Nature Neuroscience, 1.

Niv, Y., Daniel, R., Geana, A., Gershman, S. J., Leong, Y. C., Radulescu, A., & Wilson, R. C. (2015). Reinforcement learning in multidimensional environments relies on attention mechanisms. The Journal of Neuroscience, 35(21), 8145–8157.

Oemisch, M., Westendorff, S., Azimi, M., Hassani, S. A., Ardid, S., Tiesinga, P., & Womelsdorf, T. (2019). Feature-specific prediction errors and surprise across macaque fronto-striatal circuits. Nature Communications, 10(1), 176.

O’toole, A. J., Jiang, F., Abdi, H., & Haxby, J. V. (2005). Partially distributed representations of objects and faces in ventral temporal cortex. Journal of Cognitive Neuroscience, 17(4), 580–590.

Owen, S. F., Berke, J. D., & Kreitzer, A. C. (2018). Fast-spiking interneurons supply feedforward control of bursting, calcium, and plasticity for efficient learning. Cell, 172(4), 683–695.

Parker, A. J., & Newsome, W. T. (1998). Sense and the single neuron: Probing the physiology of perception. Annual Review of Neuroscience, 21(1), 227–277.

Pfeiffer, M., Nessler, B., Douglas, R. J., & Maass, W. (2010). Reward-modulated hebbian learning of decision making. Neural Computation, 22(6), 1399–1444.

Pinsk, M. A., DeSimone, K., Moore, T., Gross, C. G., & Kastner, S. (2005). Representations of faces and body parts in macaque temporal cortex: A functional MRI study. Proceedings of the National Academy of Sciences, 102(19), 6996–7001.

Poort, J., Khan, A. G., Pachitariu, M., Nemri, A., Orsolic, I., Krupic, J., Bauza, M., Sahani, M., Keller, G. B., Mrsic-Flogel, T. D., & others. (2015). Learning enhances sensory and multiple non-sensory representations in primary visual cortex. Neuron, 86(6), 1478–1490.

Price, A., Filoteo, J. V., & Maddox, W. T. (2009). Rule-based category learning in patients with Parkinson’s disease. Neuropsychologia, 47(5), 1213–1226.

Quiroga, R. Q., Reddy, L., Kreiman, G., Koch, C., & Fried, I. (2005). Invariant visual representation by single neurons in the human brain. Nature, 435(7045), 1102–1107.

Radulescu, A., Niv, Y., & Ballard, I. (2019). Holistic reinforcement learning: The role of structure and attention. Trends in Cognitive Sciences, 23(4), 278–292.

Rajan, K., Harvey, C. D., & Tank, D. W. (2016). Recurrent network models of sequence generation and memory. Neuron, 90(1), 128–142.

Ranganathan, G. N., Apostolides, P. F., Harnett, M. T., Xu, N.-L., Druckmann, S., & Magee, J. C. (2018). Active dendritic integration and mixed neocortical network representations during an adaptive sensing behavior. Nature Neuroscience, 21(11), 1583–1590.

Rissman, J., & Wagner, A. D. (2012). Distributed representations in memory: Insights from functional brain imaging. Annual Review of Psychology, 63, 101–128.

Roelfsema, P. R., & Holtmaat, A. (2018). Control of synaptic plasticity in deep cortical networks. Nature Reviews Neuroscience, 19(3), 166.

Sacramento, J., Costa, R. P., Bengio, Y., & Senn, W. (2018). Dendritic cortical microcircuits approximate the backpropagation algorithm. Advances in Neural Information Processing Systems, 8721–8732.

Small, S. L., Hart, J., Nguyen, T., & Gordon, B. (1995). Distributed representations of semantic knowledge in the brain. Brain, 118(2), 441–453.

Soltani, A., Khorsand, P., Guo, C., Farashahi, S., & Liu, J. (2016). Neural substrates of cognitive biases during probabilistic inference. Nature Communications, 7, 11393.

Song, H. F., Yang, G. R., & Wang, X.-J. (2016). Training excitatory-inhibitory recurrent neural networks for cognitive tasks: A simple and flexible framework. PLoS Computational Biology, 12(2), e1004792.

Song, H. F., Yang, G. R., & Wang, X.-J. (2017). Reward-based training of recurrent neural networks for cognitive and value-based tasks. Elife, 6, e21492.

Spitmaan, M., Horno, O., Chu, E., & Soltani, A. (2019). Combinations of low-level and high-level neural processes account for distinct patterns of context-dependent choice. PLoS Computational Biology, 15(10), e1007427.

Spitmaan, M., Seo, H., Lee, D., & Soltani, A. (2020). Multiple timescales of neural dynamics and integration of task-relevant signals across cortex. Proceedings of the National Academy of Sciences, 117(36), 22522–22531.

Sussillo, D., & Abbott, L. F. (2009). Generating coherent patterns of activity from chaotic neural networks. Neuron, 63(4), 544–57.

Taverna, S., Ilijic, E., & Surmeier, D. J. (2008). Recurrent collateral connections of striatal medium spiny neurons are disrupted in models of Parkinson’s disease. Journal of Neuroscience, 28(21), 5504–5512.

Tsao, D. Y., Freiwald, W. A., Knutsen, T. A., Mandeville, J. B., & Tootell, R. B. (2003). Faces and objects in macaque cerebral cortex. Nature Neuroscience, 6(9), 989–995.

Tyler, L. K., & Moss, H. E. (2001). Towards a distributed account of conceptual knowledge. Trends in Cognitive Sciences, 5(6), 244–252.

Wang, J., Narain, D., Hosseini, E. A., & Jazayeri, M. (2018). Flexible timing by temporal scaling of cortical responses. Nature Neuroscience, 21(1), 102–110.

Whittington, J. C., & Bogacz, R. (2019). Theories of error back-propagation in the brain. Trends in Cognitive Sciences, 23(3), 235–250.

Wilson, R. C., & Niv, Y. (2012). Inferring relevance in a changing world. Frontiers in Human Neuroscience, 5, 189.

Wolff, S. B., Gründemann, J., Tovote, P., Krabbe, S., Jacobson, G. A., Müller, C., Herry, C., Ehrlich, I., Friedrich, R. W., Letzkus, J. J., & others. (2014). Amygdala interneuron subtypes control fear learning through disinhibition. Nature, 509(7501), 453–458.

Wunderlich, K., Beierholm, U. R., Bossaerts, P., & O’Doherty, J. P. (2011). The human prefrontal cortex mediates integration of potential causes behind observed outcomes. Journal of Neurophysiology, 106(3), 1558–1569.

Yang, G. R., Joglekar, M. R., Song, H. F., Newsome, W. T., & Wang, X.-J. (2019). Task representations in neural networks trained to perform many cognitive tasks. Nature Neuroscience, 22(2), 297–306.

Zipser, D., & Andersen, R. A. (1988). A back-propagation programmed network that simulates response properties of a subset of posterior parietal neurons. Nature, 331(6158), 679–684.

